# A positive feedback loop between ZEB2 and ACSL4 regulates lipid metabolism to promote breast cancer metastasis

**DOI:** 10.1101/2023.04.03.535330

**Authors:** Jiamin Lin, Pingping Zhang, Wei Liu, Guorong Liu, Juan Zhang, Min Yan, Yuyou Duan, Na Yang

## Abstract

Lipid metabolism plays a critical role in cancer metastasis. However, the mechanisms through which metastatic genes regulate lipid metabolism remain unclear. Here, we describe a new oncogenic-metabolic feedback loop between the epithelial-mesenchymal transition (EMT) transcription factor ZEB2 and the key lipid enzyme ACSL4 (long-chain acyl-CoA synthetase 4), resulting in enhanced cellular lipid storage and fatty acid oxidation to drive breast cancer metastasis. Functionally, Depletion of ZEB2 or ACSL4 significantly reduced lipid droplets (LD) abundance and cell migration. ACSL4 overexpression rescued the invasive capabilities of the ZEB2 knockdown cells, suggesting that ACSL4 is crucial for ZEB2-mediated metastasis. Mechanistically, ZEB2 activated ACSL4 expression by directly binding to the ACSL4 promoter. ACSL4 binds to and stabilizes ZEB2 by reducing ZEB2 ubiquitination. Notably, ACSL4 not only promotes the intracellular lipogenesis and lipid droplet accumulation but also enhances fatty acid oxidation (FAO) and ATP production by upregulating the FAO rate-limiting enzyme CPT1A (carnitine palmitoyltransferase 1 isoform A). Finally, we demonstrated that ACSL4 knockdown significantly reduced metastatic lung nodes in vivo. In conclusion, we reveal a novel positive regulatory loop between ZEB2 and ACSL4, which promotes LD storage to meet the energy needs of breast cancer metastasis, and identify the ZEB2-ACSL4 signaling axis as an attractive therapeutic target for overcoming breast cancer metastasis.

**Highlights:** ZEB2 activates FAO through transcription of ACSL4

ACSL4 regulates lipid metabolism through FAO, promoting breast cancer metastasis Targeting ZEB2-ACSL4 signaling axis inhibits breast cancer metastasis

**Significance:** We describe a novel positive feedback loop between ZEB2 and ACSL4 that results in enhanced cellular lipid storage and FA oxidation to drive breast cancer metastasis.

## Introduction

More than 90% of breast cancer-related deaths are due to metastasis^1,2^. Current treatments, including endocrine therapy, chemotherapy, and radiation therapy, are ineffective in preventing breast cancer metastasis and remain the greatest clinical challenge for breast cancer treatment^3^. Cancer metastasis is associated with a process called epithelial-mesenchymal transition (EMT)^4,5^. EMT is a pre-metastatic state in which epithelial cells lose their tight junctions and convert to migratory mesenchymal cells^6^. Metastatic invasion is a highly energy-intensive process^7^. It has become increasingly recognized that metabolic reprogramming during the EMT process contributes to metastasis and tumorigenesis^8,9^. Emerging evidence suggests that lipid metabolic reprogramming plays a critical role in meeting the energy requirements of metastatic invasion^10-13^. Elucidating the mechanism by which reprogrammed lipid metabolism helps us exploit novel and attractive targets for metastatic therapeutic interventions.

Lipid metabolism includes a complex network of pathways that regulate fatty acid (FA) synthesis, storage, and degradation^14^. For lipid anabolism, FA are stored in a dynamic organelle called a lipid droplet, which is composed of a monolayer of phospholipids that covers a hydrophobic core containing neutral lipids, such as triacylglycerol (TAG) and cholesteryl esters (CE)^15^. Lipid droplets (LD) accumulation is associated with aggressiveness in many cancer types^16^, including breast^17^, brain^18^, liver^19^, lung^20^, and prostate^21,22^. Indeed, the aggregation of FA in LD is considered a priming state to prepare for metastasis^23-25^. In the case of need, FA can be released and oxidized for energy support. For example, it has been reported that fatty acids stored in lipid droplets were a crucial resource in fueling the metastatic process in pancreatic cancer ^24^. Additionally, previous studies have suggested that metastatic triple-negative breast cancer (TNBC) depends on fatty acid oxidation (FAO) to produce high ATP levels ^26,27^. Although lipid metabolism is crucial for cancer metastasis, the signalling pathway that regulates lipid metabolic reprogramming during breast cancer metastasis remains unclear.

Dysregulation of lipid metabolic enzymes has been documented in cancer metastasis^28^. Long-chain fatty acyl synthetase (ACSL) 4 belongs to the long-chain acyl-CoA synthetase ligase enzyme family ^29,30^. ACSL4 catalyses the conversion of long-chain FAs to acyl-CoAs, a necessary step for free long-chain FAs to enter the next metabolic pathway^31^. The increased expression and activity of ACSL4 have been observed in many cancer types and are well-known biomarkers of ferroptosis^32^. Although it has been reported that ACSL4 is a tumor suppressor that activates ferroptosis, many studies have suggested that ACSL4 is an oncogene that contributes to tumor progression. For example, ACSL4 promotes Hepatocellular carcinoma (HCC) cell proliferation and metastasis via lipogenesis and lipid droplet accumulation^19,33^. In prostate cancer, ACSL4 promotes cell growth, invasion, and hormonal resistance^34^. The function of ACSL4 in breast cancer has been implicated in hormone therapy resistance involving the regulation of energy-dependent transporter expression^35^. However, regulation of lipid metabolism by ACSL4 during breast cancer invasion remains unclear.

In the present study, we demonstrate a novel positive feedback loop between the EMT transcription factor ZEB2 and the essential lipid metabolic enzyme ACSL4, resulting in enhanced cellular lipid droplet accumulation and fatty acid oxidation to drive breast cancer metastasis. Mechanistically, ZEB2 activates ACSL4 expression by directly binding to the ACSL4 promoter. ACSL4 stabilizes and upregulates ZEB2 via transcriptional and post-transcriptional mechanisms. In addition, we also provide evidence that overexpression of ZEB2 or ACSL4 is associated with worse prognosis in advanced breast cancer. Our findings provide insights into lipid metabolic mechanisms during the EMT process and reveal a novel oncogenic-metabolic pathway critical for breast cancer EMT and metastasis.

## Materials and Methods

### Breast cancer cell lines and clinical specimens

Breast cancer cell lines MCF-7, MDA-MB-231, and BT549 were purchased from the American Type Culture Collection (ATCC, Beijing Zhongyuan Ltd., Beijing, China). Paclitaxel-resistant and epirubicin-resistant cell lines have been reported previously ^36,37^. All cell lines were maintained in Dulbecco’s modified Eagle’s medium (GIBCO) supplemented with 10% (v/v) FBS (BI, Biological Industries) and 1% (v/v) Pen/Strep (GIBCO) and incubated in a humidified atmosphere of 5% CO_2_ at 37 °C.

Primary breast cancer and adjacent normal tissues were collected from the volunteers at Guangzhou First People’s Hospital, Guangdong, China. This study was approved by the Ethics Committee of the Guangzhou First People’s Hospital (approval no. K2021-201-01). Fourteen patients were enrolled in this study. The specimens were then subjected to western blotting and immunohistochemistry (IHC) assays.

### Plasmid construction and cell transfection

The PLKO.1-TRC-LUC lentivirus vector containing shRNA (shACSL4-1 and shACSL4-2) and the control vector were purchased from WZ Biosciences. The virus package plasmids psPAX2 and pMD2.G were purchased from Addgene. To produce the lentivirus, the shRNA vector and package plasmids were co-transfected into 293T cells using polyethyleneimine (Polysciene). After 48 hours or 72 hours, supernatants were harvested and passed through 0.45 um filters to collect the virus and stored at −80 °C until further use. MDA-MB-231 cells were transfected with a viral solution containing 8ug/ml polybrene (Hanbio Biotechnology). 48 hours after transfection, the virus was removed from the culture and fresh medium was added. The cells were subsequently selected using 2 mg/mL puromycin to obtain a stable cell line. MCF-7 cells were transfected with HBLV-h-ACSL4-3xflag-ZsGreen-PURO or control lentiviral plasmid. The plasmids were purchased from Hanbio Biotechnology.

For the siRNA assay, 5 nM negative control siRNA or ACSL4 and ZEB2 siRNA (GenePharma) were transfected into MDA-MB-231 or paclitaxel-resistant cells using Lipofectamine 3000 (Invitrogen) according to the manufacturer’s instructions. Six hours after transfection, the medium was replaced with fresh growth medium and the cells were harvested after 24-48 hours for qPCR or western blot analysis. The siRNA sequences are listed in Supplementary Table S1.

### Western blot analysis

For western blotting, cells were lysed in WIP buffer (Beyotime) containing protease and phosphatase inhibitors. Protein levels were quantified by BCA assay (Beyotime), and equal amounts were separated using 10% SDS-PAGE electrophoresis and transferred onto a 0.22 µm PVDF membrane for probing. After blocking for 1 hour in 5% non-fat milk diluted in TBST, the membrane was incubated with various primary antibodies at 4°C overnight. The following primary antibodies were used: ACSL4 (ab155282, Abcam), ZEB2 (sc-271984, Santa Cruz), E-cadherin (3195S, CST), N-cadherin (610921, BD Biosciences), vimentin (D21H3, 5741T, CST), CPT1A (ab220789, Abcam), GAPDH (10494-1-AP, Protein Tech), Myc-Tag (71D10, 2278T, CST), and HA-Tag (C29F4, 3724T, CST). Next, secondary antibodies were added and the proteins were detected using an ECL kit (Millipore). Immunoreactive signals were visualized using the GE Amersham Imager 600 chemiluminescence system.

### Migration and invasion assay

A wound healing assay was performed to examine the migratory ability of the cells. Briefly, cells were seeded in 6-well plates at a density of 4×10^5^ cells per well in a complete medium at 37 °C. Until cells reached 80-90% density, a sterile plastic tip was used to create a wound line across the surface of the plates. The suspended cells were discarded. After The cells were cultured in reduced serum DMEM medium in a 5% CO_2_ incubator at 37 °C for 48 hours. Images were obtained using a phase-contrast microscope. Each assay was performed in triplicates.

The cell invasion capacity was measured using Matrigel-coated Transwell chambers (8.0 μm; Corning Inc.). Briefly, cells were suspended in 200 μL serum-free medium and seeded in the upper chamber of a Transwell. The lower chamber of the Transwell was filled with medium containing 10% FBS-containing medium. The Transwell chambers were then incubated at 37 °C. After culturing for 24 hours, the Transwell holes were penetrated with 4% paraformaldehyde for 20 minutes and then stained with 0.1% crystal violet solution for 15 minutes. Invading cells were imaged and counted in five random fields.

### Immunofluorescence (IF) and Immunohistochemistry (IHC) assays

For the immunofluorescence assay, cells were seeded on confocal dishes, washed in PBS three times, and fixed in 4% paraformaldehyde for 15 minutes at room temperature. The cells were blocked with 10% goat serum (AR0009, BOSTER) for 1 h at room temperature and washed three times. The cells were then incubated with the primary antibodies for 1 hour at room temperature. Subsequently, the cells were rinsed thrice with PBS and incubated with secondary antibodies for 1 hour at room temperature.

For BODIPY 493/503 and Phalloidin staining, cells were fixed in 4% PFA for 15 minutes and washed twice with PBS. The cells were then incubated with BODIPY 493/503 (1:2500 in PBS, GLPBIO) / Phalloidin (HUAYUN) for 15 minutes at room temperature. DAPI was used to stain the nuclei. Images were acquired by confocal microscopy (Ni-E-A1, Nikon, 40x) and analyzed using NIS Elements Viewer software and ImageJ.

For immunohistochemistry (IHC) assay, paraffin-embedded tissue slides were dewaxed with xylene and rehydrated using a graded series of alcohols. This was followed by antigen retrieval and blocking with 5% BSA for 60Lminutes. After that, Tissue slices were incubated with primary antibodies against ACSL4, ZEB2, Vimentin or ERα at 4 °C overnight in a humidified container and then detected with the SP Rabbit&Mouse HRP Kit (CWBIO). Images were acquired using a digital pathology scanning system (Aperio CS2).

### Metabolic analysis

For the oxygen consumption rate (OCR) assay, the XF long-chain fatty acid oxidation (LCFA) stress test Kit (103672-100) and Seahorse XF96 Analyzer were used to investigate the long-chain fatty acid OCR and extracellular acidification rate (ECAR) according to the manufacturer’s protocol.

ATP levels were determined using an ATP Assay Kit (S0026, Beyotime) according to the manufacturer’s protocol. ATP levels were calculated using luminescence signals and were normalized to protein concentrations. Glycolytic activity was determined using a Glycolysis Cell-Based Assay Kit (600450, Cayman Chemical), according to the manufacturer’s protocol.

### Untargeted lipidomics and proteomics

Cell Lipids were extracted in a chloroform-methanol mixed solution (2:1, −20 °C). The extracted cell lysates were immersed in liquid nitrogen and frozen for 5 minutes. The extracted cell lysates were placed in a 2 ml adapter, and the above steps were repeated twice. Samples were then centrifuged to pellet the proteins (5 minutes, 12000 rpm), and the supernatant was stored for analysis in a vacuum centrifugal concentrator. The sample was dissolved in 200 μl isopropanol, filtered through a 0.22 μm membrane, and detected by Liquid Chromatography-Mass Spectrometry (LC-MS). The lipidomics data were analyzed using LipidSearch software (version 4.0). The software identified intact lipid molecules based on their molecular weight and fragmentation pattern using an internal library of predicted fragment ions per lipid class. The spectra were then aligned based on retention time and the MS1 peak areas were quantified across the sample conditions. Excel 2010 was used to calculate intensity, and the R program (version 3.2.5) was used for data manipulation and statistical analyses, including unsupervised hierarchical clustering and heatmap visualization.

### Quantitative Real-time PCR

Total RNA was isolated using the TRIzol reagent (Invitrogen), and cDNAs was synthesized from total RNA using the PrimeScript™ RT Master Mix (Perfect Real Time, Takara). Quantitative real-time PCR was performed in triplicate using PowerUp SYBR Green (Thermo Fisher Scientific). The relative gene expression was measured using the 2^-ΔΔCt^ method. All primers used are listed in Supplementary Table S2.

### Luciferase reporter assays

For Luciferase assays, the 2000bp ACSL4 promoter vector containing luciferase was purchased from WZ Biosciences Inc. The −287bp, −965bp, −1038bp, and −1116bp regions of the ACSL4 promoter were cloned by PCR amplification using primers containing restriction sites MIII and Hind III. The nucleotide sequences of primers used are listed in Supplementary Table S3. 293T cells were transfected with 1ug of the-287bp, −965bp, −1038bp, −1116bp or 2000bp ACSL4 promoter luciferase reporter and an empty vector for 24 hours. After transfection, the cells were harvested and analyzed using Bright-Glo reagent (Promega) according to the manufacturer’s instructions.

### Chromatin immunoprecipitation (ChIP) assay

ChIP assays were performed using a ChIP assay kit (Cat.p2078, Beyotime Biotechnology). Briefly, MDA-MB-231 cells were grown to 90% confluence and crosslinking was performed with 1% formaldehyde for 10 minutes. Mouse anti-ZEB2 antibody or mouse IgG was used to immunoprecipitate the DNA-containing complexes. After the DNA purification (Cat. D0033; Beyotime Biotechnology), PCR, and Q-PCR were performed to detect the ZEB2-binding site in the ACSL4 promoter region. The primer sequences are listed in Supplementary Table S4.

### Co-immunoprecipitation (Co-IP) assay and Ubiquitination assay

For co-IP assays, HEK293T cells were co-transfected with ACSL4-FLAG and ZEB2-MYC plasmids. 48 hours after transfection, cells were lysed in WIP buffer (Beyotime) containing a protease inhibitor and phosphatase inhibitors for 30 minutes at 4 °C followed by centrifugation. The supernatants were immunoprecipitated with the indicated antibodies overnight at 4 °C, followed by incubation with protein A/G beads for 1 hour at 4 °C. After incubation, the beads were washed with WIP buffer and boiled in a 2×loading buffer. Protein samples were analyzed by western blotting. GST pulldown assay was performed by using GST pull down Assay Kit (FI88807, FITGENE) according to the manufacturer’s protocol. The GST control plasmid and ACSL4-GST plasmid were purchased from WZ Biosciences.

For the ubiquitination assay, HEK293T cells were transfected with HA-Ubi plasmid, ACSL4-FLAG plasmid, and ZEB2-MYC or vector plasmid. 48 hours after transfection, the cells were treated with 20 µM MG-132 for 6 hours to block the proteasomal degradation of ZEB2 before being lysed with WIP lysis buffer (Beyotime). Equal amounts of protein lysates were immunoprecipitated with anti-MYC beads and subjected to SDS-PAGE, followed by blotting with anti-HA (ubiquitin) to visualize polyubiquitinated ZEB2 protein bends.

### In vivo experiments

The Ethics Committee approved the animal experiments for Animal Experiments of the South China University of Technology. All NSG mice were purchased from the Medical Laboratory Animal Center of Guangdong Province, China. Six female NSG mice aged 6-8 weeks were used in each group for the primary tumor growth and spontaneous lung metastasis experiments. A total of 2 × 10^6^ vector control or shACSL4 cells were mixed 1:1 by volume with Matrigel (BD Biosciences). Each mouse was injected orthotopically into both the flanks. Xenograft tumor growth was measured and tumor volume was calculated as follows: volume=(length×width^2^)/2. At the experimental endpoint, mice were intraperitoneally injected with 150 mg/kg D-luciferin and imaged for 2 minutes using a live imager. The xenograft tumor and lung samples were removed and fixed to count the metastatic lung nodules.

### Statistical analysis

The results are reported as the mean ± standard error of the mean (SEM), as indicated in the figure legend. Student’s *t*-test was used for two-group comparisons. Comparisons between three or more groups were analyzed by one-way analysis for variance followed by Duncan’s test. Statistical comparisons for the LM2 lung metastasis assay were performed using the Mann-Whitney U test. All experiments with representative images, including western blotting and immunofluorescence, were repeated at least twice, and representative images are shown. P<0.05 was considered statistically significant.

## Results

### ZEB2 and ACSL4 are over-expressed and correlated in highly invasive breast cancer cells

To explore the molecular mechanism of highly invasive breast cancer, we performed RNA sequencing analysis of wild-type and two drug-resistant luminal breast cancer cell lines. Significant changes in 6155 genes was found (P < 0.05). Notably, EMT and stemness genes such as ZEB2, SNAIL, TWIST, TRAIL, Gli2, WNT, and AKT3, which are overexpressed in basal-like breast cancer (BLBC), were significantly upregulated in drug-resistant cells (Supplementary Fig. S1A). In contrast, differentiated genes such as FOXA1, ERα, E-cadherin, and GATA3, which are highly expressed in the luminal subtype, were significantly downregulated (Supplementary Fig. S1B), suggesting that drug-resistant cells underwent EMT and became stem-like cells. We noticed that the rate-limiting enzymes of fatty acid metabolism, long-chain fatty acyl synthetase 4 (ACSL4), and EMT transcription factor ZEB2 were among the top 200 upregulated (>2-fold) genes (Fig. 1A, Supplementary Fig. S2). To verify these findings in the clinical sample, we analyzed ZEB2 and ACSL4 expression in TCGA database and found that ACSL4 expression was positively correlated with ZEB2 expression (Fig. 1B, r=0.7657, p<0.001) and inversely correlated with ERα expression (Fig. 1C, r=-0.3312, p<0.001). by using the TCGA database, we compared the overall-survival (OS) between ACSL4 or ZEB2 high- and low-expression breast cancer patients. We found that patients with higher ACSL4 or ZEB2 expression, especially those with simultaneous high expression had worse prognosis than those with lower expression (Fig. 1D-1F).

**Figure 1.**
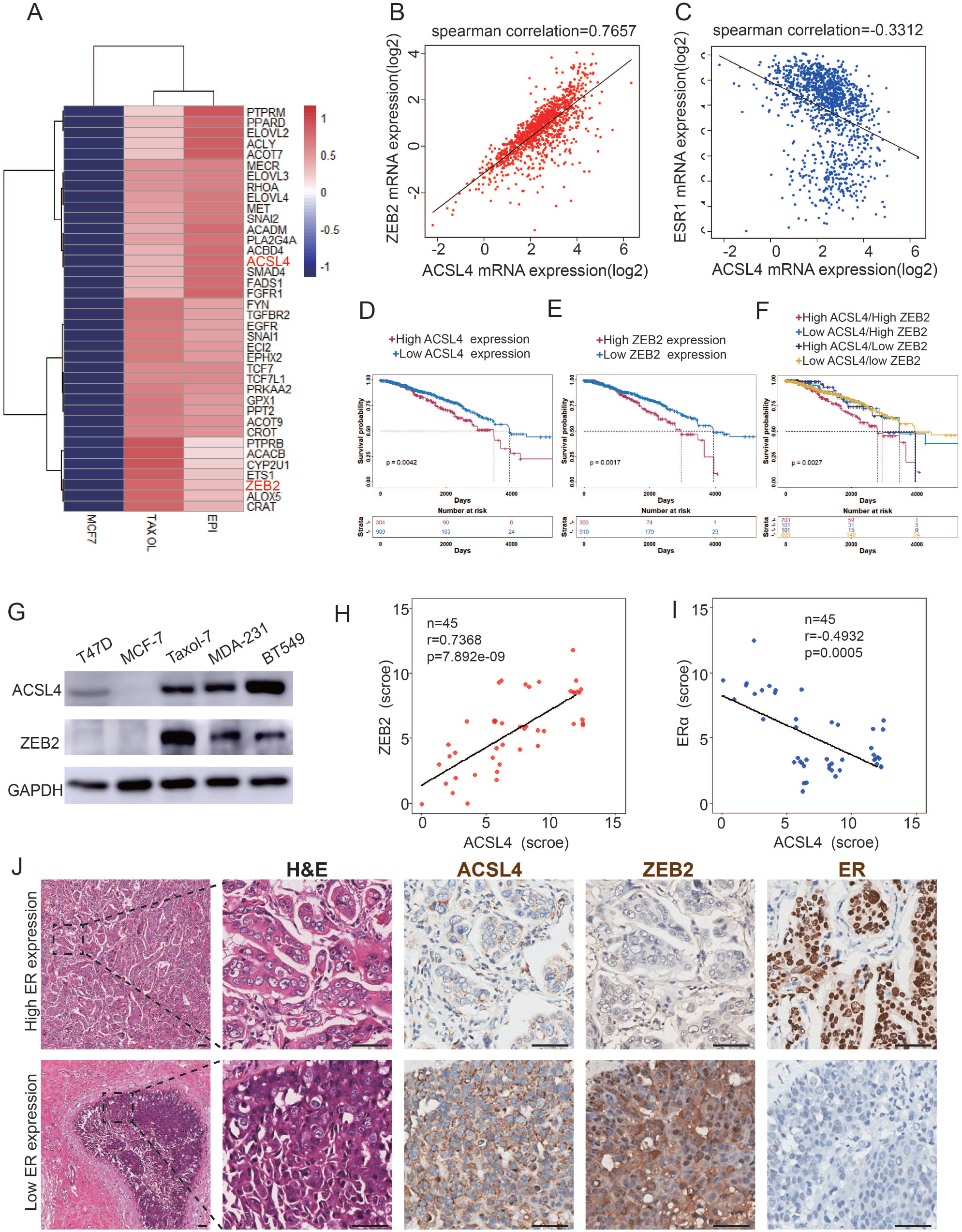
The expression and relationship of ZEB2 and ACSL4 in breast cancer. **A,** Heat maps of the 38 up-regulated EMT-related genes detected by RNA-seq analysis in the paclitaxel-resistant MCF-7 cell line (TAXOL) and Epirubicin-resistant MCF-7 cell line (EPI) compared to wild-type MCF-7 cell line. **B,** The correlation between ACSL4 and ZEB2 mRNA expression in the TCGA cohort consisting of 1222 breast cancer patient samples. Spearman correlation and linear regression analysis were employed. **C,** The correlation between ACSL4 and ER mRNA expression in the TCGA cohort consisting of 666 breast cancer patient samples. Spearman correlation and linear regression analysis were employed. **D,** OS (Overall survival) was examined by Kaplan–Meier analysis to compare the survival rates in ACSL high and low expression of breast cancer patients. **E,** OS (Overall -progression survival) as examined by Kaplan–Meier analysis to compare the survival rates in ZEB2 high and low expression of breast cancer patients. **F** OS (Overall -progression survival) was examined by Kaplan–Meier analysis to compare the survival rates in the four groups of breast cancer patients. **G,** Expression of ACSL4 and ZEB2 were analyzed by western blot in a panel of 5 breast cancer cell lines, including two basal-like (MAD-231, BT549), two luminal (T47D, MCF-7) and a taxol-resistant cell lines. **H,** The correlation between ACSL4 and ZEB2 protein expression in the IHC cohort consisting of 45 breast cancer patient samples. **I,** The correlation between ACSL4 and ER protein expression in the IHC cohort consisting of 45 breast cancer patient samples. **J,** HE staining and IHC analysis of ACSL4, ZEB2, ER expression in representative basal-like and luminal subtype breast cancer tissues. Representative pictures were shown. Scale bar, 50 μm.

To confirm this correlation, we performed western blot analysis and found that Basal-like and Taxol-resistant cell lines, which lack ERα expression, showed high expression of ZEB2 and ACSL4, whereas low or no ZEB2 and ACSL4 expression was detected in luminal subtype cell lines (Fig. 1G). Consistently, tissue samples from 45 breast cancer patients had similar expression patterns, with ZEB2 and ACSL4 being relatively highly expressed in ER-negative patient samples (patients 6, 7, 8, and 9) (Supplementary Fig. S3A and 3B). The immunohistochemistry staining assay, as shown in Fig. 1H-1J, also confirmed that the expression of ACSL4 was positively correlated with ZEB2 expression (Fig. 1H, p<0.001) and inversely correlated with ERα expression (Fig. 1I, p<0.001). Taken together, these results indicate that ACSL4 and ZEB2 are correlated and overexpressed in highly invasive breast cancers.

### Overexpression of ACSL4 contributes to ZEB2-mediated breast cancer invasion

Because ACSL4 is overexpressed in highly invasive breast cancer cells, we hypothesized that ACSL4 is essential for driving breast cancer migration and invasion. We then established a stable ACSL4 overexpressed MCF7 cell line, which was less aggressive than the basal-like breast cancer cells. The overexpression of ACSL4 significantly enhanced the metastatic and invasive capacities of MCF7 cells (Fig. 2A and 2B). Conversely, ACSL4 knockdown by shRNA significantly reduced the metastatic and invasive capacity of MDA-MB-231 cells compared to that of the control cells (Fig. 2C and 2D). Interestingly, the Phalloidin staining showed that the ACSL4 knockdown cells had a significantly smaller length to width ratio, which indicates the reversion of EMT process, than those of the control group (p<0.05)(Supplementary Fig. S4). We also observed that overexpression of ZEB2 significantly enhanced the metastatic and invasive capacities of MCF7 cells (Supplementary Fig. S5A and 4B). Conversely, the metastatic and invasive abilities of MDA-MB-231 cells were significantly reduced in ZEB2 knockdown cells (Supplementary Fig. S5C and 5D). To investigate the downstream genes regulated by ACSL4, we performed RNA sequencing analysis and identified that the tight junction and focal adhesion pathways were among the most upregulated pathways in ACSL4 knockdown cells compared to control cells (Fig. 2E). To verify the RNA-seq results, we examined the expression of adhesion-related genes by immunoblotting and found that after silencing of ACSL4 and ZEB2, the luminal epithelial marker E-cadherin was increased, whereas the mesenchymal markers, such as vimentin and N-cadherin were decreased (Fig. 2F). These results confirm the essential role of ACSL4 in breast cancer invasion and migration.

**Figure 2.**
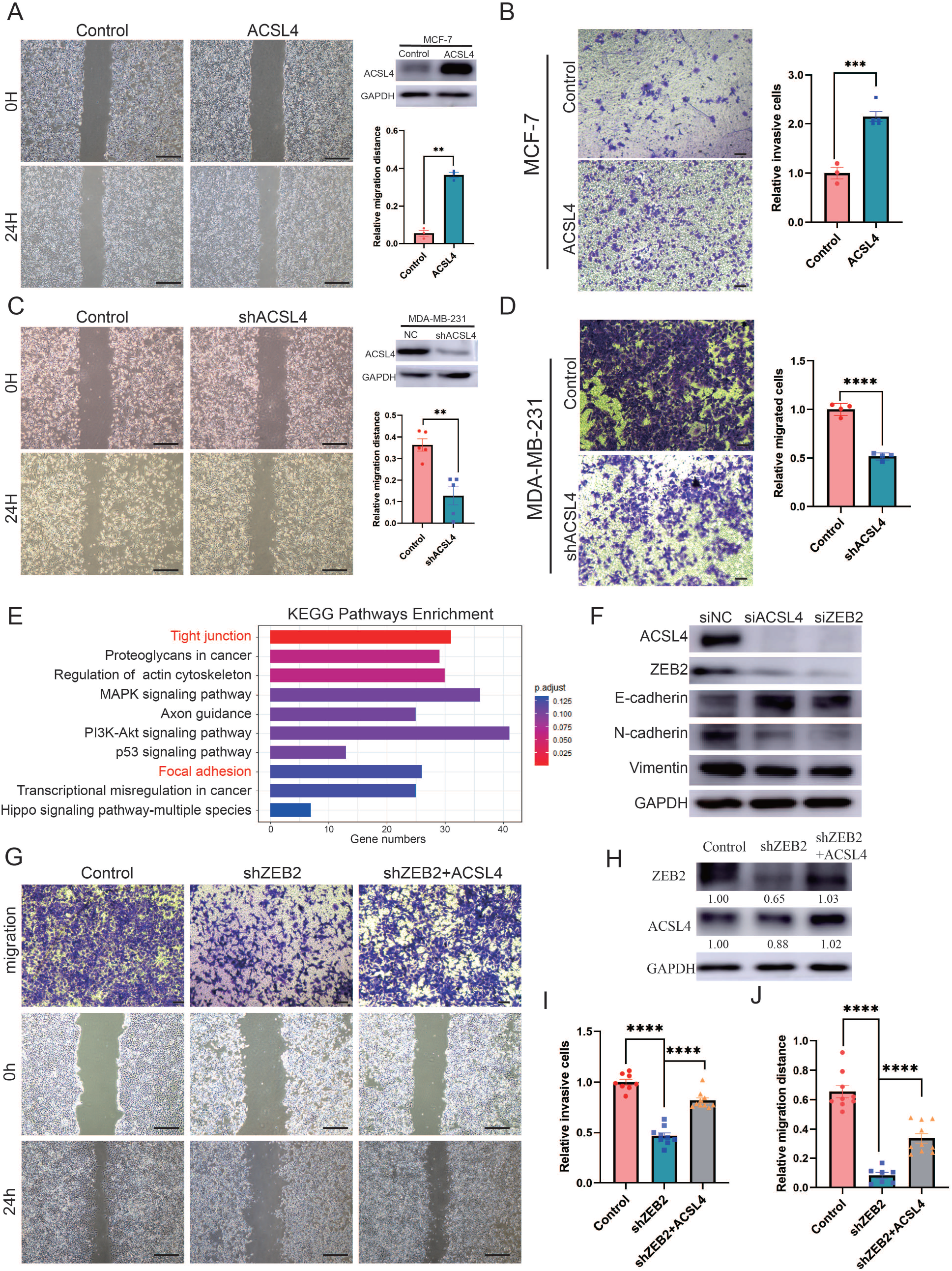
Over-expression of ACSL4 contributes to ZEB2-mediated breast cancer invasion. **A,** Cell metastatic capacity was analyzed by wound healing assay in control or ACSL4 over-expression MCF-7 cells (Left panel). Expression of ACSL4 was analyzed by western blot in control or ACSL4 over-expression MCF-7 cells. Quantification of relative migration distance (Right panel). Scale bar, 5mm. **B,** Cell invasive capacity was analyzed by transwell invasion assay in control or ACSL4 over-expression MCF-7 cells (Left panel). Quantification of relative invasive cells (Right panel). Scale bar, 1 mm. **C,** Cell metastatic capacity was analyzed by wound healing assays in control or ACSL4 silencing MDA-MD-231 cells (shACSL4) (Left panel). Expression of ACSL4 was analyzed by western blot in control or ACSL4 silencing MDA-MD-231 cells (shACSL4). Quantification of relative migration distance (Left panel). Scale bar, 5mm. **D,** Cell invasive capacity was analyzed by transwell invasion assay in control or ACSL4 silencing MDA-MD-231 cells (shACSL4) (Right panel). Quantification of relative invasive cells (Left panel). Scale bar, 1 mm. **E,** KEGG pathway enrichment analysis of differentially expressed genes by RNA-sequencing between control and ACSL4 knockdown MDA-MB-231 cells. The top 10 deferential pathways were listed. **F,** Expression of three EMT-related genes, E-cadherin, N-cadherin, and vimentin, was analyzed by western blotting in control, ACSL4, or ZEB2 silencing MDA-MB-231 cells. **G,** Cell invasive and metastatic capacity were analyzed by transwell invasion assay and wound healing assays (Scale bar, 1mm/5mm) in control and ZEB2 knockdown, or ZEB2 knockdown cells that over-express ACSL4. **H,** Protein expression was analyzed by western blot in control, ZEB2 knockdown (shZEB2), and ZEB2 knockdown with ACSL4 overexpression (shZEB2+ACSL4) cells. **I.** Quantification of relative invasive cells in G. **J,** Quantification of relative migration distance in G. Graphs indicated the Statistical analysis in G analyzed by Student’s t-test (mean ± SEM). ** P<0.01, *** P<0.001, **** P<0.0001. All results are from three or four independent experiments.

We hypothesized that the ZEB2-ACSL4 axis played a crucial role in metastatic ability. To examine whether ACSL4 was required for ZEB2-mediated breast cancer invasion and migration, ACSL4 was overexpressed in ZEB2-silencing cells (Fig. 2G-H). As expected, overexpression of ACSL4 significantly restored the invasive and metastatic capacities of ZEB2 knockdown cells by 35.1% and 32.4%, respectively (Fig. 2I and 2J), indicating that ACSL4 is essential for ZEB2-mediated breast cancer invasion and migration.

### ACSL4 and ZEB2 promote Lipid droplet production and lipogenesis

ACSL4 is a member of the long-chain acyl-CoA synthetase family, which catalyzes the conversion of long-chain fatty acids into their active forms. However, the mechanism by which ACSL4 regulates lipid metabolism in breast cancer remains unclear. A previous study revealed that ACSL4 promotes lipid droplets (LD) accumulation in hepatocellular carcinoma (HCC)^19^. Thus, we measured the basal LD content of breast cancer cells, including MCF-7, MDA-MB-231, and Taxol-resistant MCF-7 cells. The basal number and size of lipid droplets were significantly larger in MDA-MB-231 and Taxol-resistant MCF-7 cells, both of which are highly invasive breast cancer cell lines (Fig. 3A). Fluorescence microscopy revealed that ACSL4 co-localized with LD in MDA-MB-231 cells (Supplementary Fig. S6). Notably, ACSL4 knockdown reduced cytoplasmic LD abundance and LD-containing cells in MDA-MB-231 cells (Fig. 3B). Consistently, cytoplasmic LD abundance and LD-containing cells was significantly reduced in ZEB2-depleted cells (Fig. 3C).

**Figure 3.**
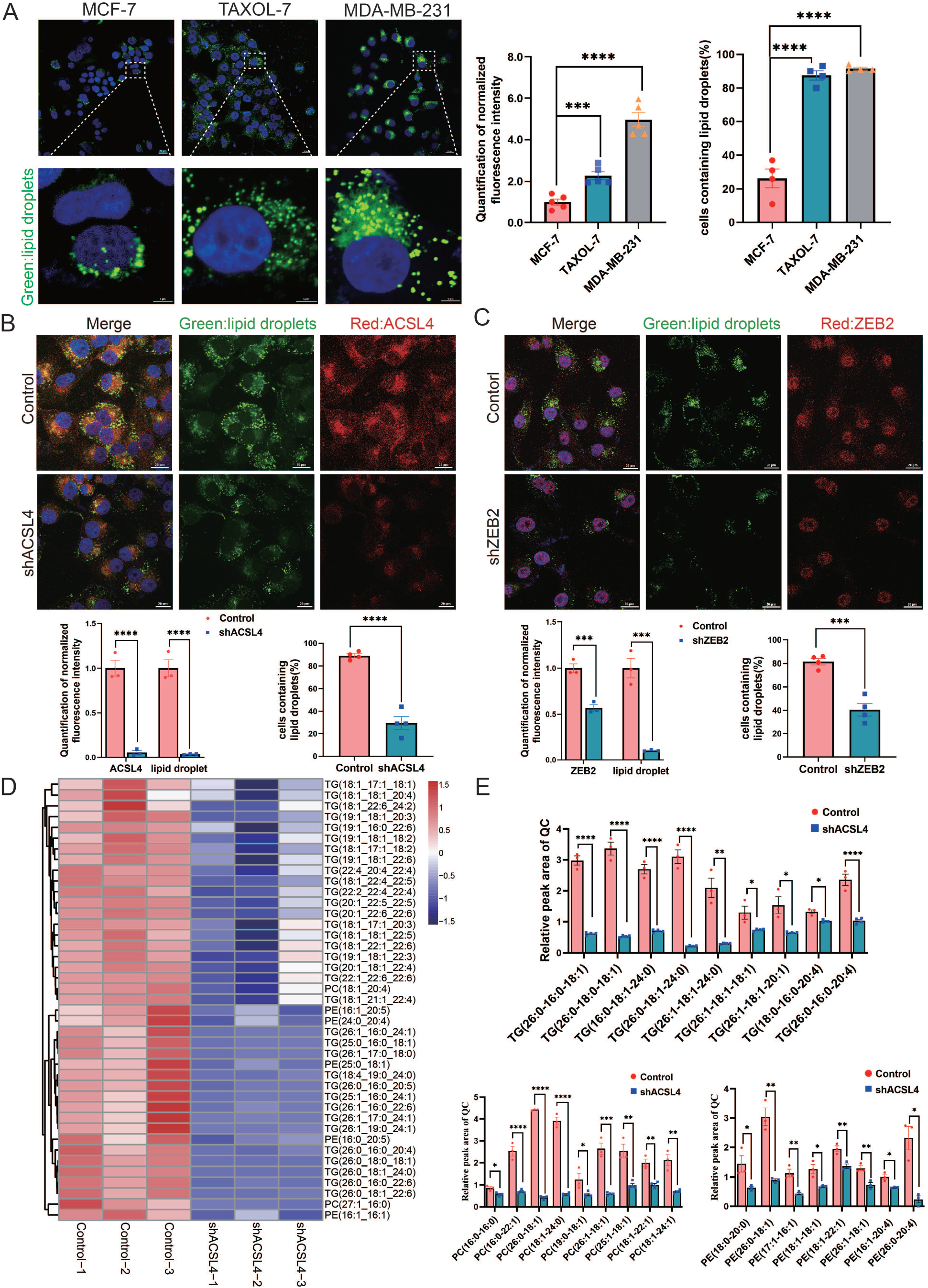
ACSL4 promotes lipid droplets accumulation and lipogenesis. **A,** BODIPY 493/503 staining of lipid droplets in MCF-7, Taxol-resistant MCF-7 cells (TAXOL-7), or MDA-MB-231 cells (Left panel). Quantification of normalized lipid contents (Right panel). Scale bar, 5μm. **B,** BODIPY 493/503 staining of lipid droplets in control or ACSL4 knockdown MDA-MB-231 cells (Upper panel). Quantification of normalized fluorescence intensity and percentage of LD-containing cell number (Lower panel). Scale bar, 50μm. **C,** BODIPY 493/503 staining of lipid droplets in control or ZEB2 knockdown MDA-MB-231 cells (Upper panel). Quantification of normalized fluorescence intensity and percentage of LD-containing cell number (Lower panel). Scale bar, 50μm. **D,** The heatmap of representative down-regulated lipid species (TG, PE, PC) with hierarchical clustering in the control cells and ACSL4 knockdown cells. Each species was normalized to the corresponding mean value, as determined by two-way ANOVA. **E,** Quantification of different Fatty Acids containing TG, PC, and PE species in control or ACSL4 knockdown MDA-MD-231 cells. * P<0.05, ** P<0.01, *** P<0.001, **** P<0.0001. Error bars, SD.

Lipid droplets are phospholipid monolayers containing a hydrophobic core comprising triacylglycerols (TG) and cholesterol esters (CE). As ACSL4 promotes intracellular LD accumulation in breast cancer cells, we investigated the effect of ACSL4 on the lipid composition of breast cancer cells. Untargeted Lipidomic analyses revealed that ACSL4 knockdown breast cancer cells had a reduced ability to incorporate long-strain monounsaturated (18:1, 17:1, 22:1, 24:1, 26:1) and saturated FA (16:0, 18:0, 24:0, 26:0) into triacylglycerol and phospholipids (PL), indicating that ACSL4 promotes the incorporation of long-chain monounsaturated and saturated fatty acids into triacylglycerol and phospholipids in these cells (Fig. 3D and 3E). Moreover, all these lipids, including triacylglycerol, phospholipids, and cholesterol esters, had decreased incorporation of several polyunsaturated FA, such as 22:6, 20:4, 22:4, and 22:5, in ACSL4 depleted cells as reported previously (Fig. 3D, Supplementary Fig. S7A)^32^. Consistently, the total amount of different lipid species, as shown in Supplementary Fig. S7B, were significantly decreased after ACSL4 knockdown. All together, these results suggest that ACSL4 directs the long train of free fatty acids into lipid anabolism to form different lipid species in the LD or other cellular membranes.

### Exogenous fatty acid promotes LD accumulation and fuel cell invasion

A previous study reported that some tumors tended to increase their intake of extracellular fatty acids to promote migration^24^. As we observed that depletion of ACSL4 greatly reduced cytoplasmic LD abundance and invasive potential in basal-like breast cancer cells, we hypothesized that LD accumulation is an important step prior to breast cancer invasion. To verify our hypothesis, we examined whether exogenous fatty acids contributed to LD accumulation and the invasive ability of breast cancer cells. Exogenous oleic acid (OA) was added to the cell culture medium. Treatment with oleic acid dramatically enhanced LD abundance in the cells (Fig. 4A), indicating that breast cancer cells tend to store lipids in LD for energy reserves. Next, we assessed the effects of exogenous oleic acid treatment on cell migration. Transwell and Wound healing assays revealed that oleic acid-treated cells exhibited significantly enhanced invasive and metastatic capacities compared with control cells. (Fig. 4B and 4C, Supplementary Fig. S8). To better determine the role of oleic acid and ACSL4 on cell migration, the oleate was added in the culture medium of ACSL4 knockdown cells. As expected, the addition of oleate obviously restores the invasive and metastatic capacities of ACSL4 knockdown cells by 33.12% and 18.61% respectively (Figure 4D).

**Figure 4.**
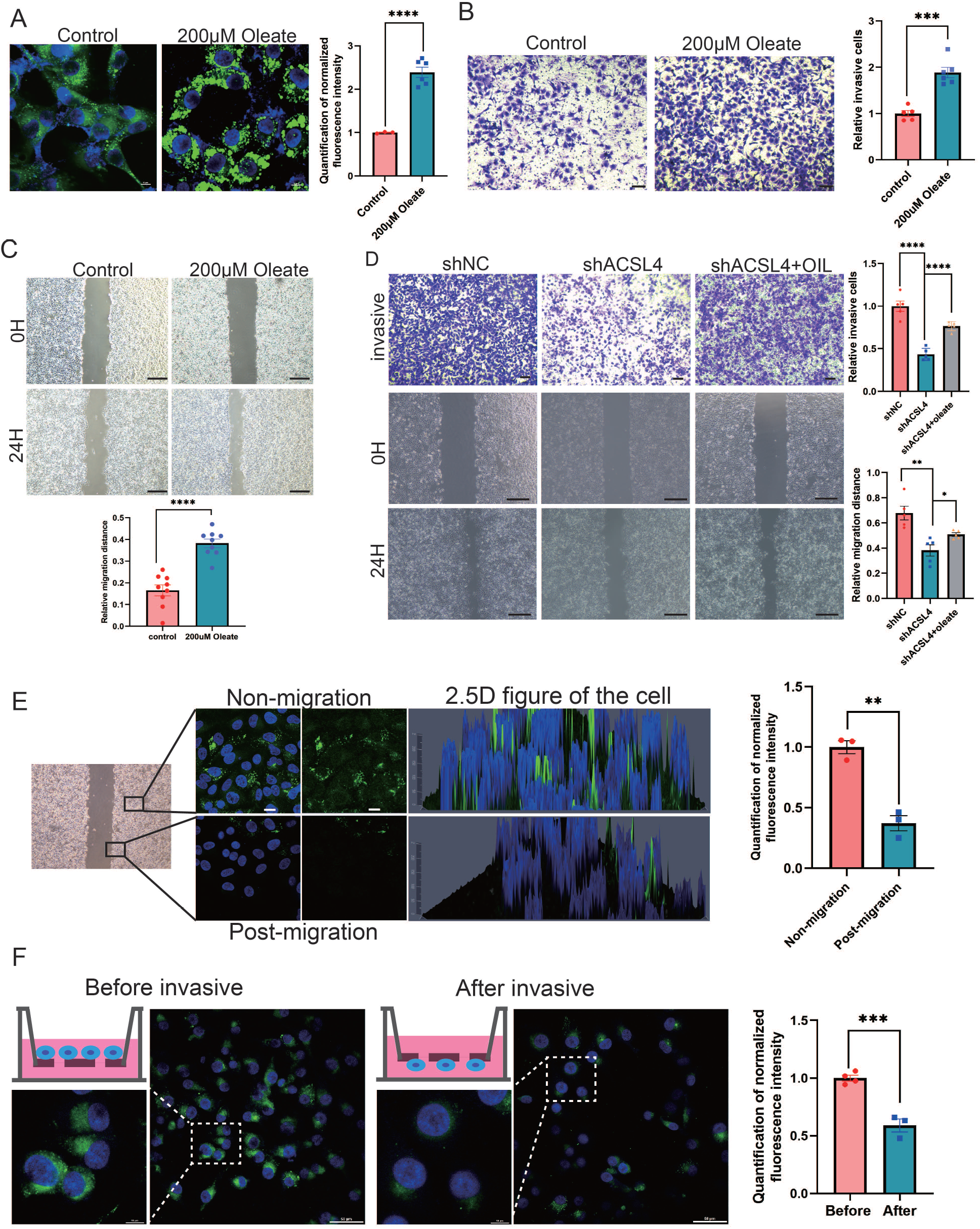
Exogenous lipids promote LD accumulation and fuel cell migration. **A,** BODIPY 493/503 staining of lipid droplets in control cells or oleic acid (OA) loaded (200 µM) MDA-MB-231 cells. Scale bar, 10 μm. Quantification of normalized lipid contents from the conditions in (A). **B,** Cell invasive capacity was analyzed by transwell invasion assay in control or oleic acid (OA) loaded (200 µM) MDA-MB-231 cells. Cells invaded for 16 hours through a Matrigel-coated filter toward high-serum media (Left panel). Quantification of relative invasive cells (Right panel). Scale bar, 1 mm. **C,** Cell metastatic capacity was analyzed by wound healing assays in control or oleic acid (OA) loaded (200 µM) MDA-MB-231 cells. Scale bar, 5 mm. **D,** Cell invasive capacity and metastatic capacity were analyzed by transwell invasion assay and wound healing assays in control, shACSL4 and oleic acid (OA) loaded ACSL4 knockdown MDA-MB-231 cells. Quantification of relative invasive cells and migrated cells. Scale bar, 1 mm /5 mm. **E,** BODIPY 493/503 staining of lipid droplets in the cells at the leading edge of the scratch and the cells that away from the edge. The 2.5 D figure of the cell was shown. Quantitation of total LD area per cell. Scale bar, 10 µm. **F,** Cells were seeded in a transwell chamber. After 24 hours, cells migrated to the lower side of the chamber, and the fluorescence intensity per cell was calculated (Left panel). Quantitation of total LD area per cell before and after cell migration (Right panel). Scale bar, 10 µm (small) and 50 µm (big). * P<0.05, ** P<0.01, *** P<0.001, **** P<0.0001.

Previous study reported that LD undergo lipolysis during the process of migration in pancreatic cancer^24^. We reasoned that breast cancer cells utilize stored lipids during migration to fuel metastasis. Lipid droplet content was analyzed using fluorescence microscopy after cell migration. We observed that the lipid signal was significantly decreasing in the leading edge of the scratch of the wound-healing migration (Fig. 4E). In addition, we observed significantly reduced LD in cells on the lower side of the transwell chamber, suggesting that lipids stored in LD were utilized and degraded during cell migration and invasion (Fig. 4F). Taken together, these results suggest that lipids stored in lipid droplets are a crucial resource to fuel the process of breast cancer invasion and migration.

### ACSL4 and ZEB2 stimulate long-chain fatty acid oxidation and ATP generation in BLBC Cells

To explore the mechanisms by which ACSL4 regulates lipid metabolism, we performed RNA sequencing to investigate downstream pathways and genes regulated by ACSL4. KEGG enrichment analysis revealed that the FAO pathway was among the top 20 regulated pathways (Fig. 5A). ACSL4 knockdown reduced the expression of genes involved in the FAO pathways (Fig. 5B). Importantly, we observed that the FAO rate-limiting enzyme CPT1A was significantly reduced in ACSL4 knockdown cells (Fig. 5B). Reverse transcription PCR confirmed that deletion of ACSL4 significantly reduced the expression of CPT1A, but did not affect the expression of CPT1B and CPT1C (Fig. 5C). Consistently, ACSL4 silencing reduced CPT1A protein expression (supplementary Fig. S9A). In addition, levels of other lipid metabolic enzymes, such as ATGL, FASN, and SREBP2, were significantly decreased after ACSL4 knockdown (Supplementary Fig. S9B). As ACSL4 knockdown decreased CPT1A expression, we reasoned that ACSL4 might stimulate fatty acid oxidation in highly invasive breast cancer. Oxygen consumption rate (OCR) was calculated in ACSL4 or ZEB2 silencing and control cells. We observed that the OCR rate derived from long-chain fatty acids was significantly reduced in ACSL4 or ZEB2 silencing cells (Fig. 5D and 5E), suggesting the metabolic advantage of increased long-chain fatty acid oxidation and oxidative phosphorylation (OXPHOS).

**Figure 5.**
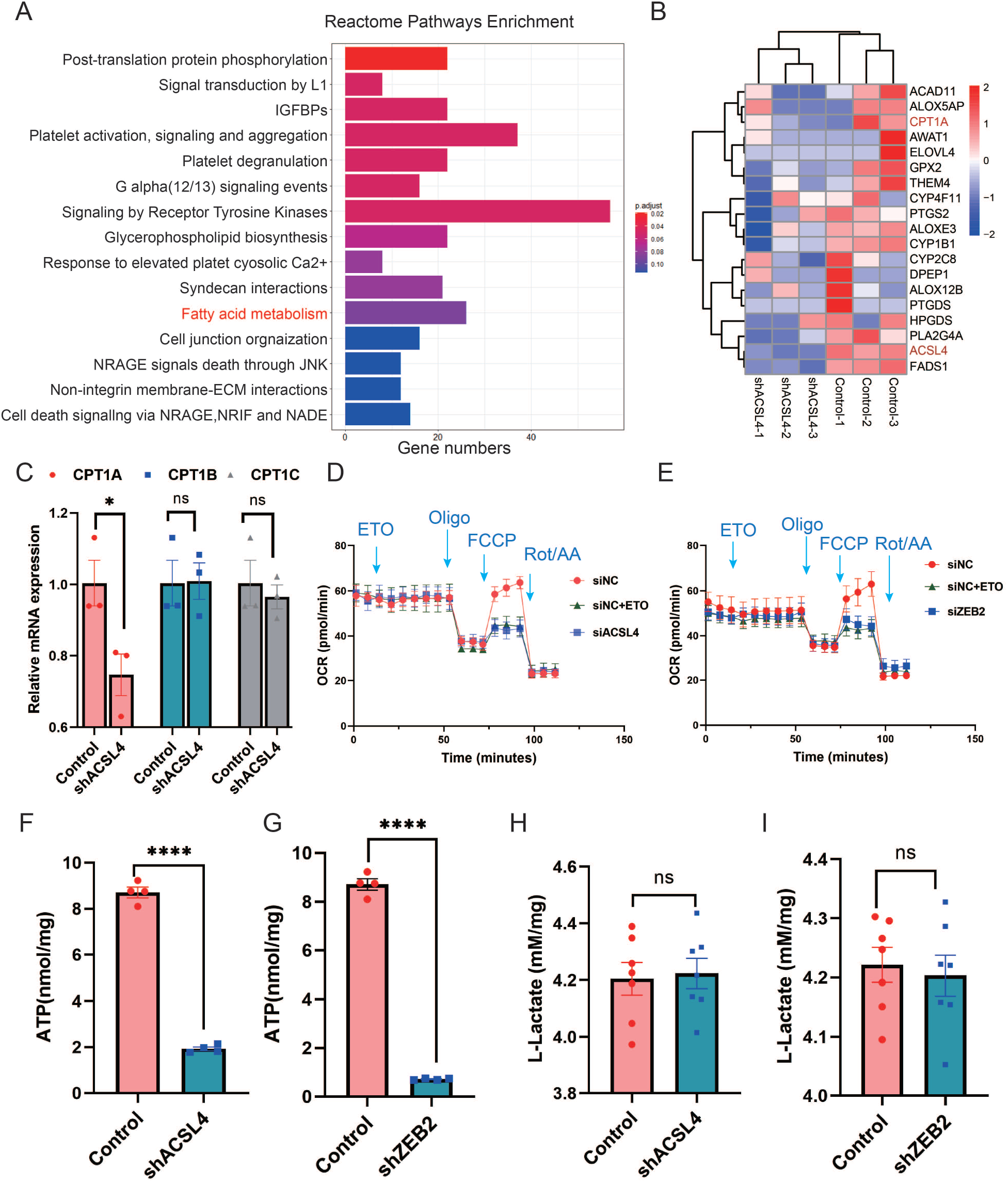
ACSL4 and ZEB2 increase fatty acid oxygen consumption and promote ATP generation in BLBC Cells. **A,** Reactome pathway analysis of differentially expressed genes by RNA-sequencing between control and ACSL4 knockdown MDA-MB-231 cells. The representative pathways were shown. **B,** Heatmap showing the differentially expressed genes between control cells and ACSL4 knockdown cells. The representative FAO-related genes were shown. **C,** The mRNA levels of CPT1A, CPT1B, and CPT1C were analyzed by quantitative PCR in control and ACSL4 knockdown cells. **D,** Quantitation of the normalized oxygen consumption rate (OCR) for long-chain fatty acids was monitored by Agilent XF Substrate Oxidation Stress Test in control or ACSL4 knockdown cells. Specific inhibitors were added as indicated. **E,** Quantitation of the normalized oxygen consumption rate (OCR) for long-chain fatty acids was monitored by Agilent XF Substrate Oxidation Stress Test in control or ZEB2 knockdown cells. Specific inhibitors were added as indicated. **F and G,** ATP production was quantified in control or ACSL4 knockdown cells (Left) or ZEB2 knockdown cells (Right). **H and I,** Lactate production was examined in control and ACSL4 knockdown (shACSL4) or ZEB2 knockdown (shZEB2) cells. Data are represented as mean SEM of three independent experiments, analyzed by Student’s t-test, * P<0.05, **** P<0.0001.

Since OXPHOS is accompanied by increased ATP generation, we measured ATP levels and found that ACSL4 or ZEB2 knockout cells had significantly reduced ATP generation compared to control cells (Fig. 5F and 5G). To exclude OXPHOS derived from the aerobic glycolysis pathway, we measured lactate production and found no significant difference in lactate production between ACSL4 or ZEB2 knockdown and control cells, suggesting that glucose metabolism is not involved in ACSL4 or ZEB2 mediated metabolic process (Fig. 5H and 5I). All these results indicated that ACSL4 upregulates CPT1A to stimulate fatty acid oxidation and ATP generation in BLBC Cells.

### ZEB2 transcriptionally activates the expression of ACSL4

ZEB2 is a crucial transcription factor involved in EMT. Because ZEB2 and ACSL4 were highly correlated in the clinical samples in TCGA database, we investigated whether ACSL4 is a direct downstream target of ZEB2. Silencing of ZEB2 by siRNA in MDA-MB-231 cells significantly reduced ACSL4 mRNA levels and protein expression in MDA-MB-231 cells (Fig. 6A and 6 B). We found that the ACSL4 promoter contained four canonical ZEB2-binding E-boxes (CAGGT/CACCT) located at −287, −965, −1038, and −1116 of the ACSL4 promoter, respectively (Fig. 6C). Therefore, we cloned five segments of the ACSL4 promoter and a control segment to generate promoter-luciferase constructs, based on the location of these E-boxes (Fig. 6C). ZEB2 overexpression significantly enhanced the luciferase activity of all five E-box-containing segments of the ACSL4 promoter, whereas no significant change was observed in the control segment (Fig. 6D). To examine whether ZEB2 directly binds to the ACSL4 promoter, we performed chromatin immunoprecipitation (ChIP) using four sets of primers (Fig. 6E). Primer set 1, which covered segment 1, consistently exhibited apparent ZEB2 binding (Fig. 6F and 6G). However, no binding was detectable using the other three sets of primers (Fig. 6F), indicating that ZEB2 binds to the E-Box located at nucleotides −184 to −295 of the ACSL4 promoter.

**Figure 6.**
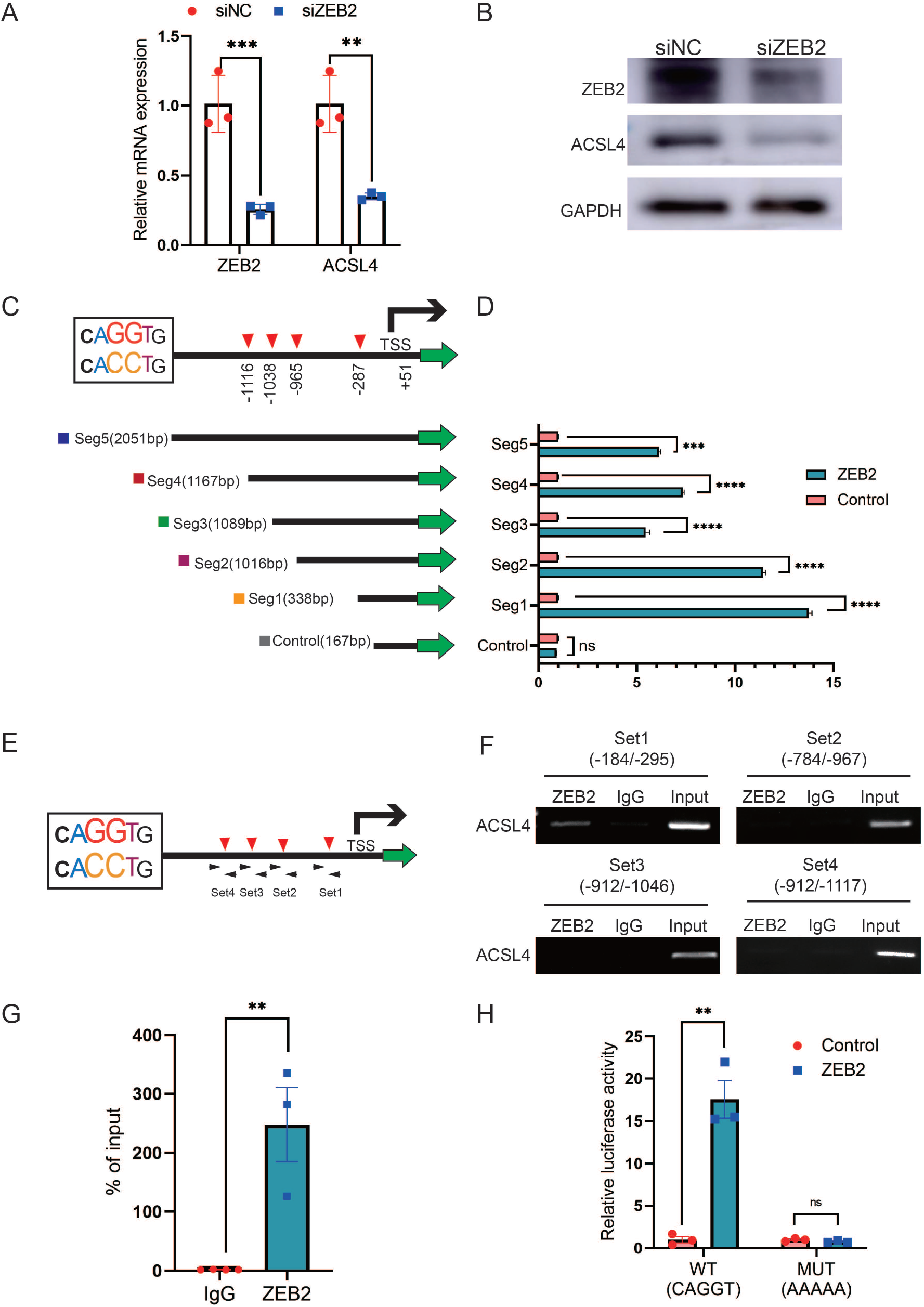
ZEB2 activates ACSL4 expression by directly binding to its promoter. **A,** Relative mRNA levels of ZEB2 and ACSL4 in control or ZEB2 knockdown MDA-MB-231 cells. **B,** Protein levels of ZEB2 and ACSL4 in control or ZEB2 knockdown MDA-MB-231 cells. **C and D,** Truncated ACSL4 promoter segment activity analyzed by Luciferase reporter assay in control or ZEB2 over-expressed 293T cells. **E,** The specific primers designed for ACSL4 promoter according to the E-box position shown in C (set1 to −287bp, set2 to −995, set3 to −1038, set4 to −1116). **F,** ChIP assay analysis of the occupation of ZEB2 on ACSL4 promoter by using four primers as indicated in E. **G,** Quantitative PCR analysis of ZEB2-binding abundance of specific ACSL4 promoter region by using set1 primer indicated in F. Genomic DNA was purified after chromatin immunoprecipitation and analyzed by Quantitative PCR. **H,** Luciferase reporter analysis of the activity of wild-type ACSL4 promoter or its mutants in control or ZEB2 over-expression 293T cells. Data are represented as mean SD of three independent experiments, analyzed by Student’s t-test, ** P<0.01, *** P<0.001, **** P<0.0001, ns: no significance.

Since the −287 E-box exhibited apparent ZEB2 binding, we generated mutants of the −287 E-box promoter-luciferase construct. The Luciferase Reporter Assay revealed that ACSL4 promoter activity was almost entirely abolished by the mutations (Fig. 6H), suggesting that this promoter region is essential for ZEB2-mediated ACSL4 promoter activation. Taken together, these data suggest that ZEB2 directly binds to the ACSL4 promoter to activate its mRNA expression.

### ACSL4 regulates ZEB2 mRNA expression and protein stabilization

We hypothesized that ACSL4 regulates the expression of ZEB2. We then performed quantitative PCR and immunoblotting and observed that both ZEB2 mRNA and protein levels were reduced after the depletion of ACSL4 in the two BCSC cell lines (Fig. 7A and 7B). A previous study has reported that ACSL4 regulates c-Myc protein stability in HCC. We reasoned that ACSL4 might in turn regulate the stability of ZEB2. Therefore, we performed a ubiquitination assay to investigate whether ACSL4 regulates ZEB2 protein stability via ubiquitination. HEK293T cells were co-transfected with HA-ubiquitin and myc-ZEB2 expression vectors along with either an empty vector or an ACSL4 overexpression vector. As shown in Fig. 7C, the expression of ACSL4 caused a significant decrease in the ubiquitination of ZEB2. Conversely, we observed a increasing ubiquitination of ZEB2 in ACSL4 silencing cells (Fig. 7D), suggesting that ACSL4 attenuated the ubiquitin proteolysis of ZEB2. Notably, co-IP assays and GST pull down assays revealed a specific interaction between ACSL4 and ZEB2 proteins, supporting the notion that ZEB2 and ACSL4 are present in a protein complex (Fig. 7E and 7F, Supplementary Fig. S10). Immunofluorescence assay revealed that ACSL4 and ZEB2 were co-localized in some certain regions of the cytoplasm (Supplementary Fig. S11). To further confirm the role of ACSL4 in the regulation of ZEB2 proteolysis, cycloheximide was added to the cell medium to block the synthesis of new proteins. Interestingly, endogenous ZEB2 protein levels were almost completely suppressed in ACSL4 knockdown cells from 0h to 8h time points (Fig. 7G and 7H). This phenomenon could be explained by the fact that ZEB2 mRNA levels were significantly suppressed by the ACSL4 knockdown. Conversely, ZEB2 protein maintained a relatively steady level at 6 hours in ACSL4 overexpressing cells compared with control cells that undergone obvious ZEB2 proteolysis at 4 hours after adding CHX, indicating an increased ZEB2 protein stability in ACSL4 overexpressing cells (Fig. 7I and 7J). Taken together, these results indicate that ACSL4 not only stabilizes the ZEB2 protein by attenuating its ubiquitination but also upregulates ZEB2 mRNA expression. Therefore, ACSL4 regulates ZEB2 through both transcriptional and post-transcriptional mechanisms.

**Figure 7.**
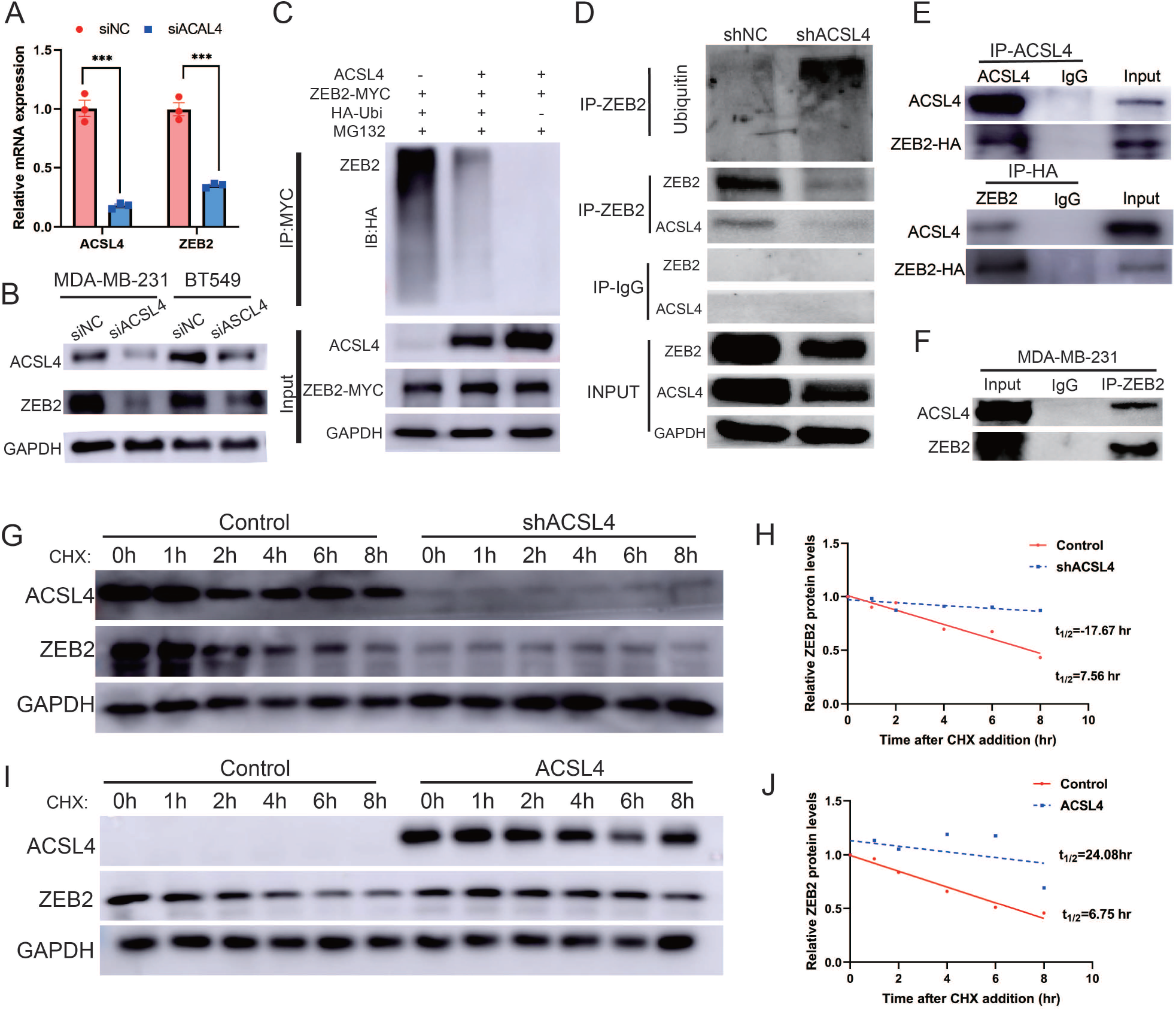
ACSL4 regulates ZEB2 mRNA expression and protein stability. **A,** Relative mRNA levels of ZEB2 and ACSL4 in control or ACSL4 silencing MDA-MB-231 cells. **B,** Protein levels of ZEB2 and ACSL4 in control or ACSL4 silencing MDA-MB-231 cells as indicated. **C,** Ubiquitylation of ZEB2 was examined by in vitro ubiquitin assay. 293T cells were co-transfected with indicated constructs. Cells were treated with MG132 for 6 hours before IP. Anti-MYC was used to pull down the ZEB2 protein. The polyubiquitinated ZEB2 protein was detected by an anti-HA antibody. **D,** Ubiquitylation of ZEB2 was examined in MDA-MB-231 cells. The indicated antibody was used to pull down the protein in control or ACSL4 knockdown MDA-MB-231 cells. The polyubiquitinated ZEB2 protein was detected by anti-ubiquitin antibody. **E,** The interaction between ACSL4 and ZEB2 was detected by Co-IP assay. 293T cells were co-transfected with ZEB2 and ACSL4 expressing construct.The indicated antibody was used to pull down the protein. **F,** The interaction between ACSL4 and ZEB2 was detected by IP assay in MDA-MB-231 cells. Anti-ZEB2 antibody was used to pull down the protein. **G,** The stability of ZEB2 protein was detected by CHX treatment assay in control or ACSL4 silencing MDA-MB-231 cells. Cells were treated with 100 µg/mL cycloheximide (CHX) and were harvested at the indicated times after the addition of CHX. GAPDH was used as the internal loading control. **H,** Quantification of stability assays shown in G. **I,** The stability of ZEB2 protein was detected by CHX treatment assay in control or ACSL4 over-expressed MCF-7 cells. GAPDH was used as the internal loading control. **J,** Quantification of stability assays shown in I. *** P<0.001.

### ACSL4 knockdown inhibits lung metastasis of BLBC in the animal model

To further validate our in vitro findings, we examined the effect of ACSL4 on lung metastasis of breast cancer cells in an animal model. Depletion of ACSL4 resulted in a significant reduction in tumor growth and lung colonization of highly metastatic MDA-MB-231 cells (Fig. 8A). In contrast, the control group showed significantly more lung metastatic nodules (Fig. 8B) and rapid tumor growth (Fig. 8C and 8D). Immunohistochemistry confirmed a striking downregulation of ACSL4, accompanied by ZEB2 and vimentin, compared to those in the control group (Fig. 8E).

**Figure 8.**
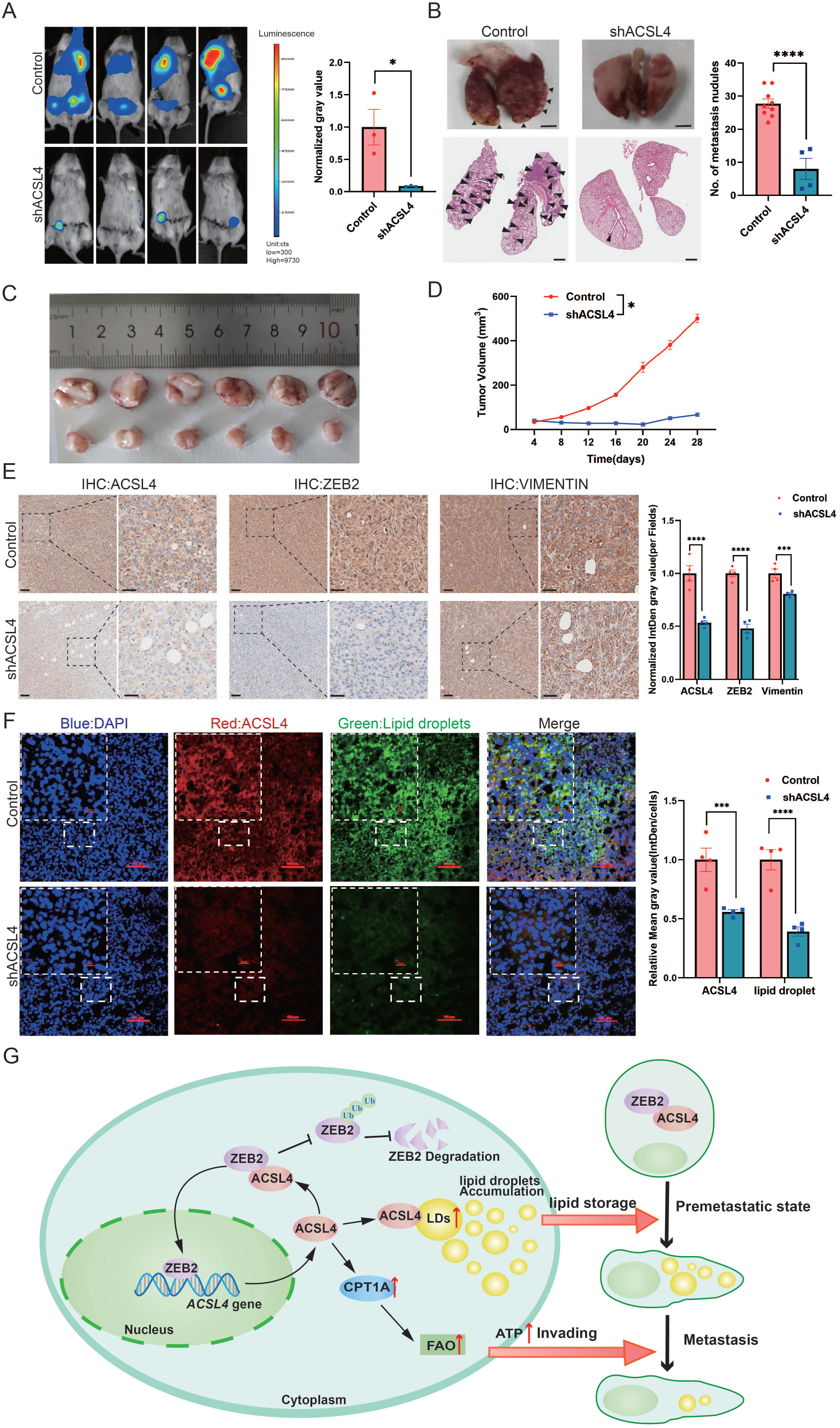
ACSL4 knockdown attenuates lung metastasis of breast cancer. **A,** Cell metastatic capacity was determined by xenograft experiment in vivo. Control cells and ACSL4 knockdown MDA-MB-231 cells were mixed with Matrigel in a 1:1 ratio and injected into NSG mice. Lung metastatic burden in mice was quantified by bioluminescence imaging at the experimental endpoint (Left panel). Quantification of normalized gray area (Right panel). **B,** Representative xenograft tumor images and HE staining pictures of metastatic nodules in lungs (pointed by black arrows) were shown. The right panel shows the quantification of metastatic lung nodules in two groups as indicated. **C,** The image of xenograft tumors developed from the control and ACSL4 knockdown cells. **D,** The tumor volumes were measured and calculated at the indicated time. **E,** Representative IHC images of lung metastatic lesions (Left panel) and quantification is shown (Right panel). Scale. Bar, 50μm. **F,** The representative images of fluorescence assay for ACSL4 and lipid droplets (Left panel). The fluorescence intensity of lipid droplets and ACSL4 were calculated in the right panel. Scale. Bar, 100μm. **G,** A proposed mechanism to illustrate the positive feedback loop of ZEB2 and ACSL4 regulates lipid metabolism, which results in enhanced breast cancer metastasis. Data are represented as mean SEM of three independent experiments and analyzed by Student’s t-test, * P<0.05, *** P<0.001, **** P<0.0001.

To further validate the role of LD in ACSL4 high-expression cells, we examined LD content in frozen sections of the two groups. Consistent with the in vitro results, there was an obvious reduction in LD accumulation in the ACSL4 depletion tissue compared to that in control tissue (Fig. 8F). Taken together, these data suggest that ACSL4 promotes the lung metastasis of breast cancer cells in vivo.

## Discussion

The rewiring of metabolic pathways during EMT has only recently been recognized. In addition to glycolysis, dysregulated lipid metabolism has been shown to contribute to cancer invasion and metastasis^38,39^. Many EMT-driving factors have been found to reprogram the lipid metabolic pathways by regulating metabolic enzymes. For example, TGF-β, a major driver of EMT, activates fatty acid synthase (FASN) and forms a FASN-TGFβ1-FASN positive loop in cisplatin-resistant cells, resulting in EMT induction^40-42^. However, the direct regulation of lipid enzymes by EMT factors remains to be elucidated. Herein, we demonstrated that the lipid rating-enzyme ACSL4 is a direct downstream target of the EMT transcription factor ZEB2 in controlling lipid metabolism. Mechanistically, ZEB2 activates ACSL4 mRNA expression by directly binding to its promoter, which contains four ZEB2 consensus sequences. Importantly, we observed a strong correlation between ZEB2 and ACSL4 expression levels in clinical breast cancer samples. ACSL4 re-expression rescues the migration ability of ZEB2-depleted cells. These observations indicated that ACSL4 is crucial for ZEB2-mediated metastasis.

We also demonstrated that ACSL4 directly binds to and stabilizes ZEB2 by reducing its ubiquitination. Our results are consistent with a recent report that ACSL4 stabilizes the oncoprotein c-Myc via the ubiquitin-proteasome system^33^. We propose that as an LD enzyme, ACSL4 may participate in the protein degradation system of LD. This is likely to be a novel function of ACSL4. Interestingly, a recent study reported that lipid droplets recruit Numb through the AP2A/ACSL3 complex to promote Numb degradation^43^. Thus, it is likely that lipid droplets act as a platform for protein degradation. Interestingly, our RNA-seq data revealed that some ubiquitin E3 ligases, such as FBXO4, UBE3C, NEDD4, RBX1 etc. were significantly reduced in ACSL4 knockdown cells (Supplementary Fig. S12). This result indicated that ACSL4 may reduce the ubiquitin of ZEB2 via down-regulating ubiquitin E3 ligase. Additionally, we found that ACSL4 promoted ZEB transcription as the mRNA level of ZEB2 was significantly reduced after ACSL4 knockdown. A recent study reported that LD-derived lipolysis provide acetyl-CoA for the epigenetic regulation of gene transcription^44^. We observed that ACSL4 can also promote FAO, which generates acetyl-CoA for the epigenetic regulation. It is likely that ACSL4 regulates the ZEB2 mRNA level via lipid-epigenetic reprogramming mechanism, which is worth studying in the future. Therefore, ACSL4 regulates ZEB2 not only via a post-transcriptional mechanism but also via a transcriptional mechanism. Notably, the relationship between ZEB2 and ACSL4 could form a positive feedback loop: ZEB2 transcriptionally activates ACSL4, and conversely, ACSL4 stabilizes the ZEB2 protein by reducing its ubiquitination. Amplification of either gene might therefore lock this loop in an active state, resulting in the enhanced invasive and metastatic capabilities of breast cancer cells (Fig. 8G).

Previous studies have reported that ACSL4 could act as a tumor suppressor or oncogene, depending on the specific cancer type and tissue environment^45-49^. Indeed, ACSL4 could either be located at the cytomembrane or at the lipid droplets and endoplasmic reticulum membrane, indicating the different functions of ACSL4^30^. The cytomembrane ACSL4 is likely responsible for dictating ferroptosis sensitivity by shaping the plasma membrane lipidome, whereas ACSL4 localized to lipid droplets has other functions. We provide evidence that ACSL4, which is located in lipid droplets, is a pro-metastatic factor that promotes invasion and migration. Notably, we demonstrated that ACSL4 depletion significantly suppressed the invasion and migration of breast cancer cells in vitro and in vivo. Furthermore, ACSL4 and ZEB2 were found to be preferentially expressed in basal-like breast cancer cells and clinical samples that lacked ERα expression, indicating that they are promising targets for highly invasive breast cancer. Survival analysis revealed that breast cancer patients with high expression of ACSL4 or ZEB2 is associated with worse overall survival than those patients with low expression. Notably, patients with high expression of both have the worst overall survival. Thus, ACSL4 and ZEB2 could specifically be used as prognostic markers for breast cancer, and this axes hold promise as a new metabolic therapeutic target for highly invasive breast cancer.

Although ACSL4 has been shown to play an essential role in metastasis in many types of cancer ^34,50,51^, its mechanism of ACSL4-mediated metastasis is not fully understood. ACSL4 is a member of the long-chain fatty acetyl-CoA synthetase enzyme family that catalyzes fatty acids to their active form, acyl-CoA, which can be directed to anabolism or catabolism, depending on the cellular background. We observed that the knockdown of ACSL4 significantly reduced the number and size of LD, indicating that fatty acids were directed to anabolism to form LD by ACSL4. Indeed, it has been shown that fatty acids are stored in LD before entering fatty acid oxidation, and the formation of LD is required for fatty acids oxidation ^11,23,52^. This is in line with our observation that exogenous fatty acids (oleic acid) significantly enhance lipid droplet accumulation and migration. Importantly, lipidomic analysis revealed that the knockdown of ACSL4 significantly decreased the incorporation of both saturated and unsaturated fatty acids into different species of lipids, including triacylglycerols (TAG), phospholipids (PE, PC), and cholesteryl esters (CE). Consistently, Frozen tissue sections from a metastatic animal model confirmed that high ACSL4 expression was accompanied by apparent LD accumulation. These results suggest that ACSL4 is a crucial regulator that promotes lipid storage during breast cancer metastasis. Interestingly, knockdown of ACSL4 restored the expression of E-cadherin, an essential adhesion molecule, indicating that ACSL4 promotes tumor invasion through multiple mechanisms.

Reprogramming of lipid metabolism is an essential step in metastasis. Lipid droplets are highly dynamic monolayer membrane-bound organelles involved in energy utilization, signal transduction, and cancer invasion. The number and size of LD are associated with cancer aggressiveness^15^. We observed that highly invasive breast cancer cells were enriched with LD. Using fluorescence microscopy, we noticed that the number of LD was significantly reduced after cell migration. Our findings are in line with a recent study showing that LD undergo lipolysis during the process of migration in pancreatic cancer^24^. It is likely that fatty acids are released from lipid droplets and form acyl-CoA, which enters OXPHOS to support the energy needed for metastasis. Therefore, these data support the notion that increased LD accumulation is a priming state prior to metastasis, and that LDs are crucial resources to fuel the process of metastasis.

In addition, we observed that ACSL4 also participates in FA catabolism, as the long-chain fatty acid-derived OCR rate and ATP production were significantly reduced in ACSL4-depleted cells, indicating that ACSL4 is essential for FAO stimulation. Our results are in line with a recent study that reported that hexokinase 2 enhances tumorigenicity by activating the ACSL4-mediated fatty acid β-oxidation pathway ^53^. Notably, the expression of CPT1, a rate-limiting enzyme of FAO, was significantly reduced in the ACSL4-depleted cells. Among the three isoforms of CPT1, CPT1A is the only isoform regulated by ACSL4. We proposed that ACSL4 promotes FAO and ATP production by upregulating CPT1A, thereby providing energy support for breast cancer metastasis. Our results reveal the mechanism of previous findings that FAO is a critical energy pathway in triple-negative breast cancer (TNBC)^26^. In addition to the finding that ACSL4 promotes FAO to fuel metastasis, we observed that lactate levels did not change after ACSL4 knockdown, suggesting that glycolysis is not involved in ACSL4-mediated energy metabolism. Therefore, we propose a model in which cell migration is a multistep process accompanied by dynamic cellular lipid metabolic changes. At the pre-metastatic stage, ACSL4 enhances LD accumulation by promoting lipogenesis. During metastasis, ACSL4 stimulates fatty acid oxidation to generate adenosine triphosphate (ATP).

In conclusion, we provide insights into the mechanistic links between EMT and lipid metabolism and identify the ZEB2/ACSL4 axis as a novel metastatic metabolic pathway that stimulates both lipogenesis and fatty acid oxidation, resulting in enhanced breast cancer invasion and metastasis. Importantly, our results demonstrate that ACSL4 is a direct downstream target of ZEB2 in controlling lipid storage and LD accumulation, which are important steps and energy pools for metastasis. Clinically, our findings identified ZEB2-ACSL4 signalling as an attractive therapeutic target for overcoming breast cancer metastasis. Elevated ACSL4 levels can be used as an effective marker for predicting cancer progression in patients with advanced breast cancer. The limitation of this study is the clinical samples is only 45. The future study should expand the clinical samples and cases to provide more clinical evidence for the crucial role of ACSL4 in breast cancer metastasis.

## Acknowledgements

This research was supported by the Guangzhou Science and Technology Project (202201020240 to N. Yang), the Medical Research Foundation of Guangdong Province (A2021112 to N. Yang), the National Natural Science Foundation of China (81602434 to N. Yang), the Research Agreement between South China University of Technology and Guangzhou First People’s Hospital (D9194290 to Y. Duan), and the National Natural Science Foundation of China (81972594 to M. Yan). We acknowledge Professors Quentin Liu for the paclitaxel-resistant and epirubicin-resistant MCF-7 cell lines and Dr. Jue Wang and Dr. Peilin Liao for their technical support. We thank all members of Professor YY. Duan’s lab for helpful insights and valuable advice.

## Authors’ Contributions

N. Yang and J. Lin designed the study. J. Lin, P. Zhang, J. Zhang, and W. Liu performed experiments. W. Liu and G. Liu provided the clinical samples. N. Yang, J. Lin, and P. Zhang analyzed the data. N. Yang, J. Lin, M. Yan, and Y. Duan wrote and revised the manuscript. N. Yang, M. Yan and Y. Duan provided financial support.

## Disclosure of Potential Conflicts of Interest

The authors declare no potential conflicts of interest.

## Supplementary Figure Legends

**Supplementary Figure S1:** The representative differential genes between wild-type MCF-7 cells, paclitaxel-resistant MCF-7 cells (TAXOL), and Epirubicin-resistant MCF-7 cells (EPI). **A,** The heatmap of the up-regulated genes in drug-resistant cells was shown. **B,** The heatmap of the down-regulated genes in the drug-resistant cell was shown.

**Supplementary Figure S2:** The volcano Plot was generated using R4.3.0 software for differentially expressed genes in the paclitaxel-resistant MCF-7 cell line (TAXOL) and Epirubicin-resistant MCF-7 cell line (EPI) compared to wild-type MCF-7 cell line.

**Supplementary Figure S3: A,** Expression of ACSL4 and ZEB2 were analyzed by western blot in a panel of nine cases of tumour samples, including 5 cases of luminal (Patient 1, 2, 3, 4, 5) and 4 cases of triple-negative breast cancer (Patient 6, 7, 8, 9). **B,** Scatter plot with linear regression analysis from expression level of A.

**Supplementary Figure S4:** Fluorescence staining of Phalloidin in control or ACSL4 knockdown MDA-MB-231 cells. The morphology of cells was analyzed by the ratio of length to width (Right panel). *** P<0.001.

**Supplementary Figure S5:** ZEB2 increases the metastatic and invasive capacities in breast cancer cells. **A,** Cell metastatic capacity was analyzed by wound healing assay in control or ZEB2 over-expression MCF-7 cells (Left panel). Quantification of relative migration distance (Right panel). Scale bar, 5mm. **B,** Cell invasive capacity was analyzed by transwell invasion assay in control or ZEB2 over-expression MCF-7 cells (Left panel). Quantification of relative invasive cells (Right panel). Scale bar, 1mm. **C,** Wound healing assays were performed in control cells or ZEB2 knockdown (shZEB2) MDA-MD-231 cells (Left panel). Quantification of relative migration distance (Right panel). Scale bar, 5 mm. **D,** Transwell invasive assay was performed in control cells or ZEB2 knockdown (shZEB2) MDA-MD-231 cells (Left panel). Quantification of relative invasive cells (Right panel). Scale bar, 1mm. ** P<0.01, *** P<0.001, **** P<0.0001.

**Supplementary Figure S6:** Fluorescence staining of lipid droplets and ACSL4 in MDA-MB-231 cells. Scale bar, 10 µm.

**Supplementary Figure S7:** ACSL4 regulates lipid composition in BLBC cells. **A,** The heatmap of the down-regulated lipid species in ACSL4 knockdown cells (shACSL4) compared with those in control cells. **B,** Quantification of different species of lipids in two groups as indicated. * P<0.05, ** P<0.01.

**Supplementary Figure S8:** Oleate induces cell migration in breast cancer cells. Cell invasive and metastatic capacity were analyzed by transwell invasion assay and wound healing assay in control or oleic acid (OA) loaded (200 μM) MCF-7 cells. Quantification of relative invasive cells (Right panel). Scale bar, 1 mm (small), 5 mm (big). * P<0.05.

**Supplementary Figure S9:** ACSL4 regulates the expression of lipid metabolic genes. **A.** Western blot analysis of CPT1A expression in control and ACSL4 knockdown, or ZEB2 knockdown MDA-MD-231 cells. **B,** Relative mRNA levels of ATGL, FASN and SREBP2 in control or ACSL4 knockdown MDA-MB-231 cells (shACSL4). ** P<0.01, *** P<0.001.

**Supplementary Figure S10:** GST-pull-down assay was used to examine the interaction between ACSL4 and ZEB2 in MDA-MB-231 cells.

**Supplementary Figure S11:** Fluorescence staining of ACSL4 and ZEB2 in MDA-MB-231 cells. Scale bar, 10 µm.

**Supplementary Figure S12:** The differentially expressed genes between control cells and ACSL4 knockdown cells were shown. The representative ubiquitination related proteolysis pathway genes were shown.

## Notes

### Competing Interest Statement

The authors have declared no competing interest.

### Summary of Updates

Dr Min Yan is a co-corresponding author,we have submit a revised version to bioRxiv. We have also corrected the description error of "siACSL4" into "shACSL4" in the Figure S4.

