## supplementary for "A positive feedback loop between ZEB2 and ACSL4 regulates lipid metabolism to promote breast cancer metastasis"

### Supplementary information

#### Supplementary Tables and Figures

**Supplementary Table S1.** The sequences of siRNA target

| Gene name | | sequence |
|---|---|---|
| ACSL4 | sense（5'-3'） | GAGCGATTTGAAATTCCAA |
| ACSL4 | antisense（5'-3'） | TTGGAATTTCAAATCGCTC |
| ZEB2 | sense（5'-3'） | GCACAUCAGCAGCAAGAAATT |
| ZEB2 | antisense（5'-3'） | UUUCUUGCUGCUGAUGUGCTT |

**Supplementary Table S2.** The sequences of gene-specific primers used for qRT-PCR

| Gene name | | sequence |
|---|---|---|
| ACSL4 | F-Primer | CATCCCTGGAGCAGATACTCT |
| ACSL4 | R-Primer | TCACTTAGGATTTCCCTGGTCC |
| ZEB2 | F-Primer | AATGCACAGAGTGTGGCAAGGC |
| ZEB2 | R-Primer | CTGCTGATGTGCGAACTGTAGG |
| GAPDH | F-Primer | GGAGCGAGATCCCTCCAAAAT |
| GAPDH | R-Primer | GGCTGTTGTCATACTTCTCATGG |
| CPT1A | F-Primer | GATCCTGGACAATACCTCGGAG |
| CPT1A | R-Primer | CTCCACAGCATCAAGAGACTGC |
| CPT1B | F-Primer | TGTATCGCCGTAAACTGGACCG |
| CPT1B | R-Primer | TGTCTGAGAGGTGCTGTAGCAC |
| CPT1C | F-Primer | TGCCATGTCGTTCCATTCTCCC |
| CPT1C | R-Primer | GCCGACTCATAAGTCAGGCAGA |

**Supplementary Table S3.** The control/-287bp/-965bp/-1036bp/-1116bp and -2000bp regions and motif1 sequences of primers used for the ACSL4 promoter vector constructs.

| Gene name | | sequence |
|---|---|---|
| seq1 | F-Primer | GATAGGTACCGAGCTCTTACGCGTGCGA GCGGGGGCG |
| seq2 | F-Primer | GATAGGTACCGAGCTCTTACGCGTCAGG TGAGGGCGTGGG |

| | | |
|---|---|---|
| seq3 | F-Primer | GATAGGTACCGAGCTCTTACGCGTTCAG GTGGTAAGGCATTTTATATATACATATA TATATACACACACACAA |
| seq4 | F-Primer | GATAGGTACCGAGCTCTTACGCGTTCCA GGTACCTACATTTCAACAAGCAC |
| seq5 | F-Primer | GATAGGTACCGAGCTCTTACGCGTTTTA GTGTCACCTGGGCTGCTTATTAAAAATTCA |
| | R-Primer | AGTACCGGAATGCCAAGCTTCCGGAATG CCAAGCTTACTTAGA |
| mofit1 | F-Primer | TACGCGTAAAAAGAGGGCGTGGGCCAAT TCTGCGCCT |
| | R-Primer | CGCCCTCTTTTTACGCGTAAGAGCTCGGT ACCTATCG |

**Supplementary Table S4.** The sequences of gene-specific primers used for ChIP assay

| Gene name | | sequence |
|---|---|---|
| Set1-184bp/-295 | F-Primer | TCACCTGGGCTGCTTATT |
| | R-Primer | GTGTGCATCACAATTATCTGGG |
| set2-784/-967bp | F-Primer | CTCCAGGTACCTACATTTCAA |
| | R-Primer | TTGTGCTTGTGTGTGTGTATATATATATG |
| set3-912/-1046bp | F-Primer | CTCAGGTGGTAAGGCATTT |
| | R-Primer | CCCAAAAAAATAAATCTCAAGAATTCTTCCA |
| set4—912/-1117bp | F-Primer | GCAAGCCGCAGGTGAGGGC |
| | R-Primer | GATCCGCTTCTGTCAGTCTCGCTGC |

#### Supplementary Fig. S1

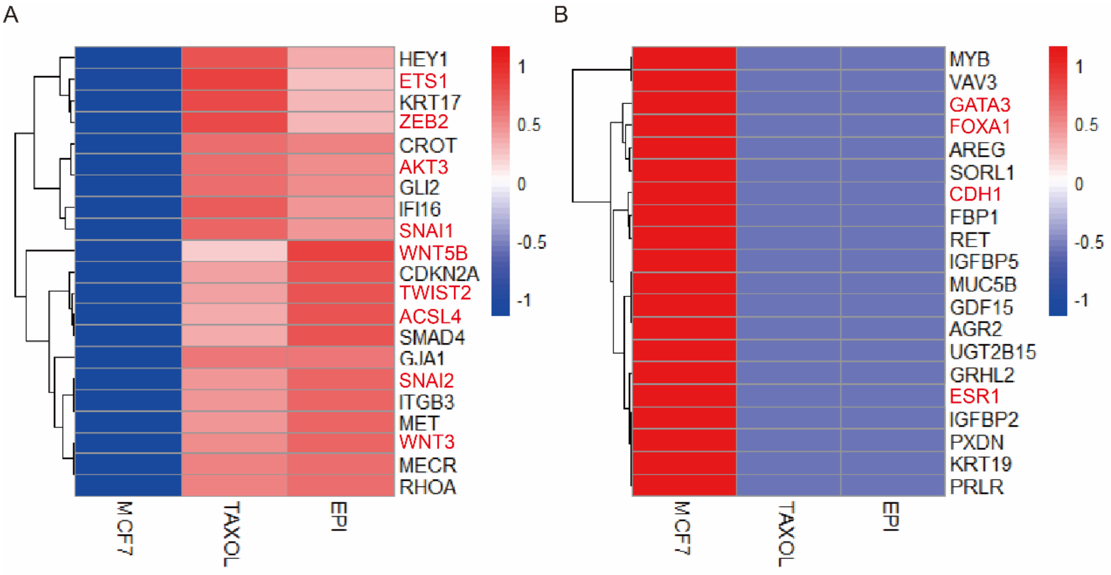


#### Supplementary Fig. S2

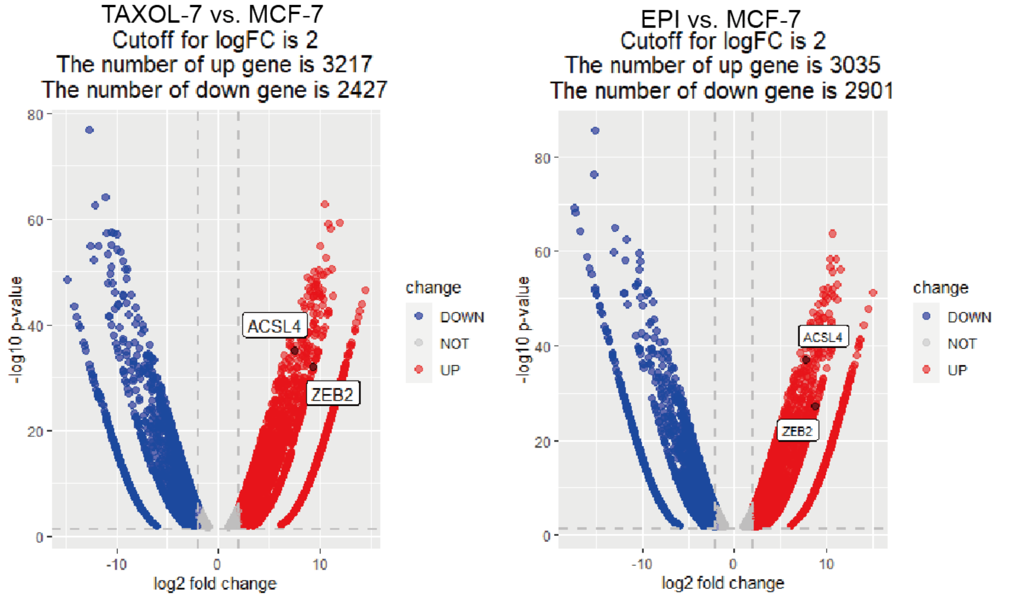

#### Supplementary Fig. S3

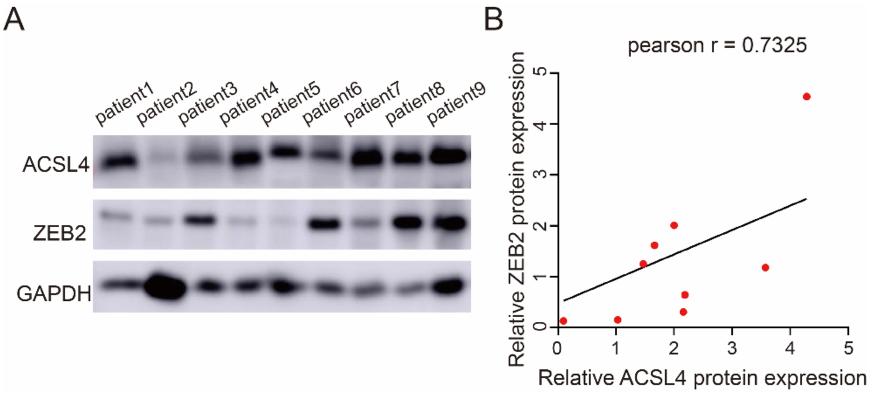


#### Supplementary Fig. S4

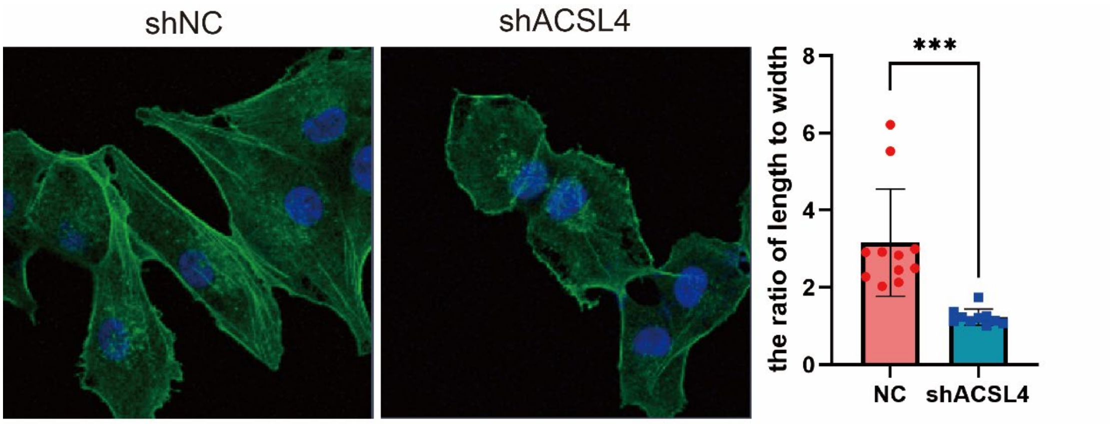

#### Supplementary Fig. S5

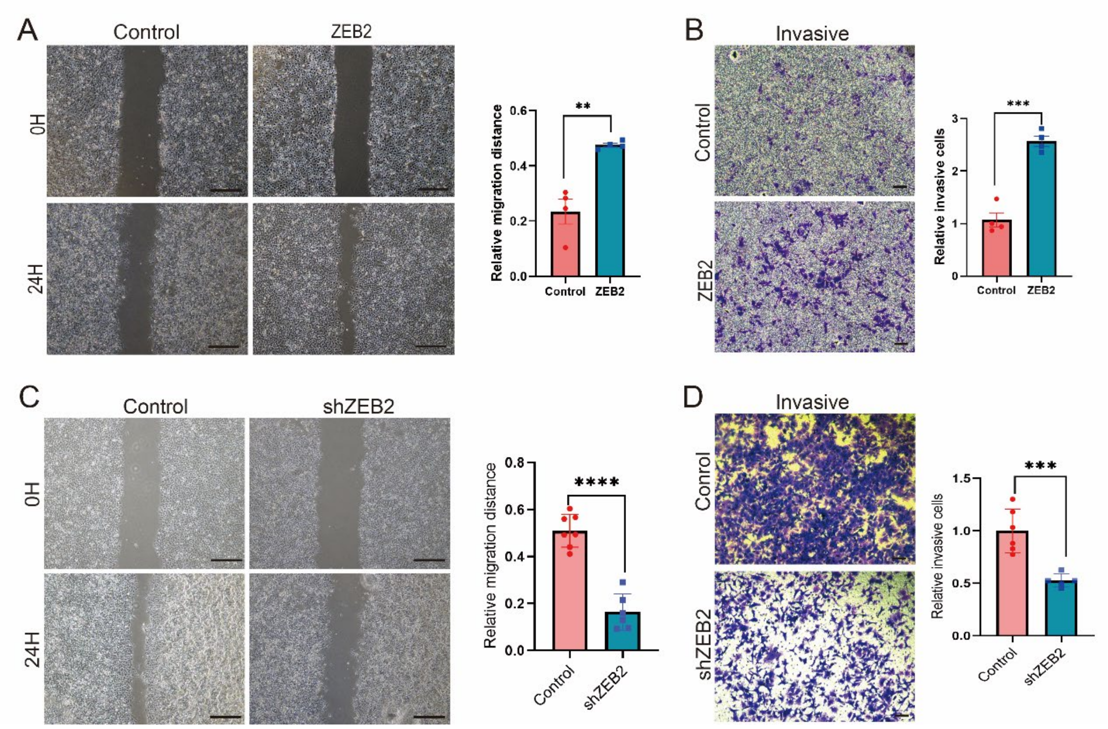


#### Supplementary Fig. S6

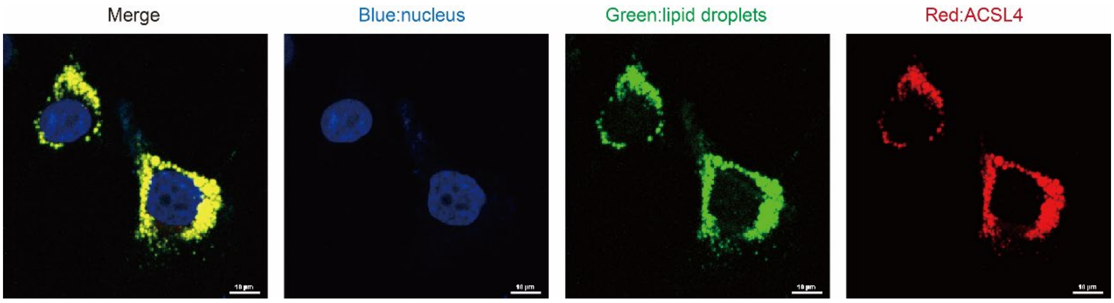

#### Supplementary Fig. S7

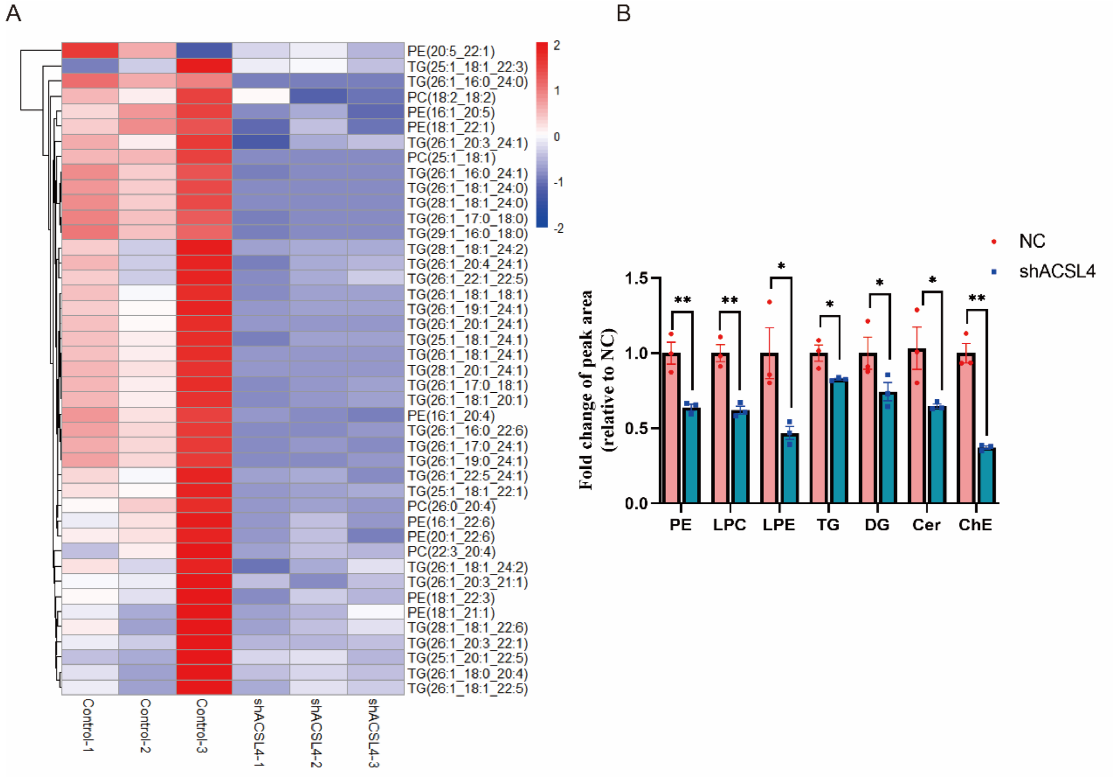


#### Supplementary Fig. S8

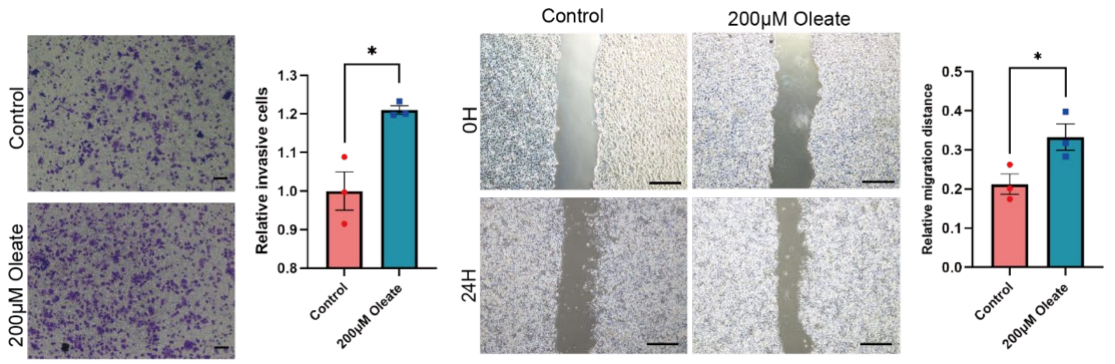


#### Supplementary Fig. S9

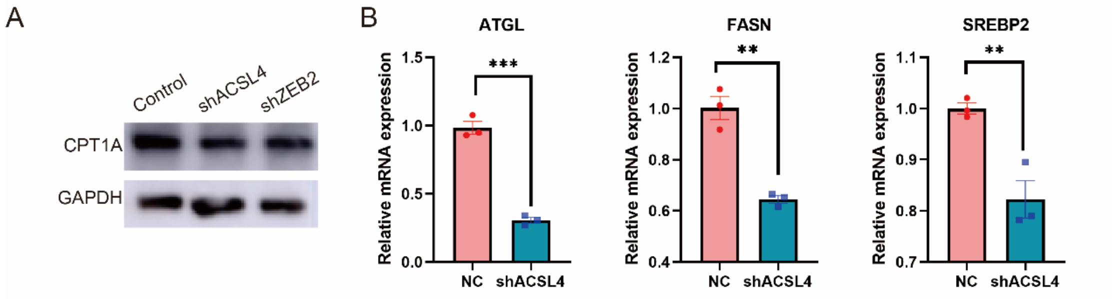

#### Supplementary Fig. S10

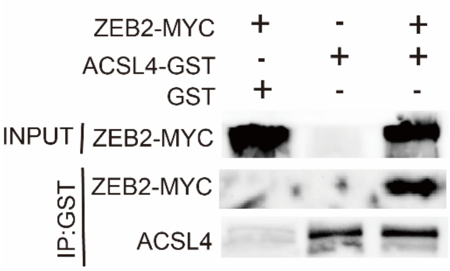


#### Supplementary Fig. S11

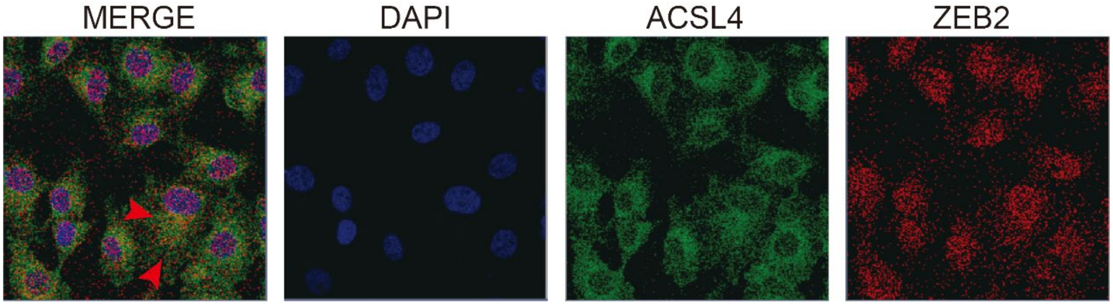


#### Supplementary Fig. S12

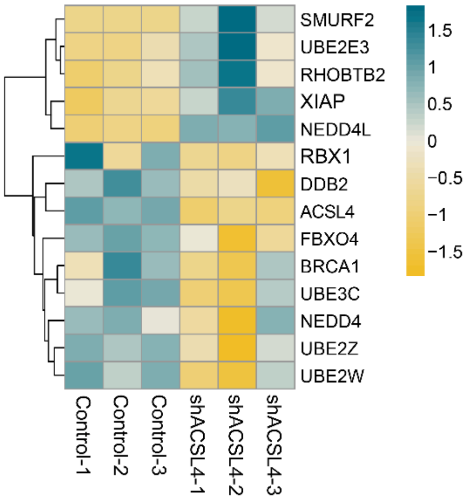
